# Validation of behavioural tests for personality assessment in tie-stall dairy cows

**DOI:** 10.64898/2026.09.25.754475

**Authors:** Xu (Aimee) Xu, Marjorie Cellier, Nadège Aigueperse, Abdoulaye Baniré Diallo, Elsa Vasseur

**Affiliations:** Department of Animal Science, McGill University, Sainte-Anne-de-Bellevue, Quebec, Canada; Université Clermont Auvergne, INRAE, VetAgro Sup, UMR Herbivores, F-63122 Saint Genès-Champanelle, France; Département d’informatique, Université du Québec à Montréal, Montréal, Québec, Canada; Research and Innovation Chair in Animal Welfare and Artificial Intelligence (WELL-E), McGill University, Sainte-Anne-de-Bellevue, Quebec H9X 3V9, Canada

**Keywords:** Animal personality, dairy cattle, behavioural testing, personality traits, welfare, personality validation

## Abstract

The study of animal personality has gained increasing attention in recent years, as researchers recognize that non-human animals exhibit stable individual differences in behaviour. In dairy cattle, understanding personality traits holds potential for improving animal welfare, enhancing management practices, and supporting productivity outcomes. However, validating the behavioural tests used to assess these traits remains a significant challenge. The present study aimed to evaluate the consistency and validity of five commonly used behavioural tests in dairy cows: the Human Approach Test (HAT), Sudden Test (SUD), Novel Object Test (NOT), Open Field Test (OFT), and Novel Surface Test (NST). These tests were chosen for their potential to assess personality dimensions that parallel components of human personality, particularly Openness, Neuroticism, Extraversion, and Agreeableness. Repeated assessments were conducted across three time periods to evaluate both intra-test (temporal) and inter-test (contextual) consistency. Results revealed that only the SUD exhibited significant behavioural consistency across all periods, suggesting its potential as a reliable tool for personality assessment. Additionally, a moderate correlation between the HAT and SUD was observed, indicating that these tests may capture overlapping personality traits. In contrast, the remaining tests (NOT, OFT, NST) lacked consistent behavioural patterns over time and across contexts, raising questions about their reliability and the complexity of trait expression. These findings highlight the challenges of identifying stable personality traits in cattle and emphasize the need for more rigorous validation of behavioural tests. This research contributes to the growing field of animal personality by providing insights into trait stability, the limitations of current testing methods, and the potential application of personality assessment in future livestock management practices.

## 1. INTRODUCTION

In recent years, the study of personality has expanded beyond humans, sparking growing interest in how non-human animals exhibit unique, stable traits. Personality, traditionally understood in human psychology, refers to the consistent patterns of affect, cognition, and behaviour that characterize individuals and distinguish them from others across time and situations (Pervin & John, 1997). A personality trait refers to a consistent response to a particular stimulus, with individuals within a population differing in the extent to which they express that response (e.g., boldness or sociability) (Woodrum Setser et al., 2023). To identify these traits, psychologists often rely on self-reported questionnaires or behavioural observations, tools designed to reveal the differences that distinguish individuals (Adolphs & Xu, 2024). This broader interest has led to a surge in research on personality in non-human animals, reflecting increased acceptance of personality constructs beyond humans (Pastorino et al., 2017). Non-human animals, like humans, exhibit stable traits that persist over time. However, identifying personality in animals relies primarily on patterns of observable behaviour. This challenge likely stems from the fact that animals, unlike humans, cannot verbally express their emotions or experiences, making observable behaviours the primary means of assessing personality (Adolphs & Xu, 2024).

While this approach avoids the challenges posed by differences in language and communication abilities, it also raises the question of whether personality is being measured in the same way across species. Since human personality assessments often rely on self-reporting, applying similar frameworks or methods to animals requires careful consideration to avoid anthropomorphic assumptions while still capturing meaningful individual differences.

Additionally, behaviours, personality traits, and overarching personality dimensions exist at different levels of analysis, where behaviours reflect traits, and traits contribute to broader personality constructs. This interdependence means that inaccuracies in measuring one level can influence the interpretation of the others (Kaiser & Müller, 2021). Furthermore, no universally accepted method exists for identifying personality traits in animals, as not all behaviours directly contribute to personality. For instance, a basic feeding behaviour may have no relevance to boldness (Kaiser & Müller, 2021), highlighting the challenge of distinguishing which behaviours are truly indicative of personality traits and useful for personality assessments.

To study animal personality, researchers often use frameworks that group specific traits, typically inferred from behaviours reflecting underlying characteristics or tendencies, into broader personality dimensions. These dimensions represent larger sets of traits that encompass various behaviours and tendencies, collectively forming an individual’s overall personality profile (Kaiser & Müller, 2021). As a result, each individual is expected to display a unique combination of these dimensions, which together define their distinct personality. Gosling and John (1999) adapted the human Five-Factor Model (McCrae & John, 1992) to propose a parallel of the five personality dimensions in animals: Openness, Extraversion, Neuroticism, Agreeableness, and Conscientiousness.

However, Conscientiousness has been inconsistently observed across species and has primarily been documented in chimpanzees (Gosling & John, 1999), while the remaining four dimensions are comparatively more widely applied in animal personality research. This inconsistency could be due, in part, to the challenges of applying human-based definitions of traits, such as organization and reliability in Conscientiousness (Roberts et al., 2014), which are typically self-reported in humans, to non-human species, where behavioural observations are the primary means of assessment and might not capture the full scope of the traits encompassed as they are defined in humans.

These limitations are particularly evident in dairy cow research, where the validation of behavioural tests used to assess personality traits remains limited. The challenge lies in interpreting behavioural expressions in a way that accurately reflects stable traits while avoiding anthropomorphism. As a result, reliable and valid methods for evaluating personality in cattle remain underdeveloped. While some studies have begun addressing this gap (Foris et al., 2018; Neave et al., 2020; Hasenpusch et al., 2024), no comprehensive review has yet synthesized an in-depth analysis of personality traits and related behavioural responses in this context (Vazire et al., 2007). A recent scoping review (Xu et al. 2026) identified only 34 relevant articles, revealing a relatively fragmented body of research with inconsistent methodologies and limited integration of trait-based frameworks. Notably, only a few studies linked observed behaviours to clearly defined personality dimensions or evaluated the long-term stability of traits, highlighting the urgent need for methodological refinement and validation in this area of study.

Yet, despite these challenges, recognizing and assessing personality in dairy cattle remains essential, not only for its implications for welfare, health, and performance, but also for its practical value in informing management strategies tailored to individual behavioural tendencies. A growing body of research highlights the relevance of personality to animal welfare and performance outcomes in farm animals (Koolhaas & Van Reenen, 2016; Marino & Allen, 2017). For instance, exploratory dairy calves have been shown to consume more solid feed at earlier ages when weaned at a fixed time point (Neave et al., 2018), a behaviour associated with improved growth performance. Calmer cows, by contrast, have been found to display stronger fear responses and lower productivity when transitioning from conventional to automated milking systems (Morales-Piñeyrúa et al., 2023). These findings demonstrate the economic and welfare benefits of incorporating personality assessments into livestock management. Beyond performance metrics, personality research also provides valuable insights into stress-related health outcomes (Mehta & Gosling, 2008; Koolhaas, 2008), aligning with growing ethical concerns around farm animal welfare.

Building on this, personality-based management strategies may offer a practical way to support herd-level welfare. Research indicates that cows with similar personalities are more likely to form strong social bonds, characterized by increased affiliative behaviour, closer proximity, and fewer aggressive interactions (Morton et al., 2015; Schuett et al., 2011). This underscores the potential of personality assessments to inform socially compatible grouping and improve overall welfare outcomes. In production environments, where competition for resources can intensify social stress, identifying and grouping compatible individuals may help reduce conflict and foster positive social relationships. This aligns with previous findings by Val-Laillet et al. (2009), who demonstrated that increased competition at the feeder reduced affiliative behaviours like allogrooming, particularly in vulnerable individuals such as primiparous or low-ranking cows. Incorporating personality assessments into grouping strategies could therefore offer a practical means of reducing social stress and improving welfare in intensive management systems.

However, for such strategies to be effective, the assessments themselves must be reliable. To bridge the gap between theory and application, researchers rely on the assumption that while behavioural responses may vary depending on test conditions, traits that are accurately identified and linked to specific behaviours should remain stable across contexts and over time, particularly when compared to others within the same group (Réale et al., 2010; Kaiser & Müller, 2021). Assessing behavioural consistency over time not only provides insights into an individual’s personality but also helps validate the tests themselves: if measured behaviours remain stable, then the test is a reliable tool for personality assessment. Thorough validation is therefore essential to ensure that commonly used behavioural tests accurately capture stable individual differences in dairy cows.

This study evaluates dairy cow personality using five well-established tests in research: the Human Approach Test (HAT), which measures responses to an unfamiliar human; the Sudden Test (SUD), which assesses reactivity to an unexpected stimulus; the Novel Object Test (NOT), where the animal’s reaction to the exposure of an unfamiliar object is quantified; the Open Field Test (OFT), which examines exploration in an unfamiliar open space and reactions to isolation; and the Novel Surface Test (NST), which assesses a cow’s response to an unfamiliar texture underfoot while balancing neophobia and social motivation, as they must cross it to rejoin their companions. These tests have been widely used to capture key personality traits such as sociability, fearfulness, and exploratory tendencies (Woodrum Setser et al., 2023; Górecka-Bruzda et al., 2011). However, there is currently a lack of consensus on how consistently these tests measure stable traits across time and context, due in part to methodological variation and the absence of short and long-term validation. Repeated testing further complicates the process, as animals may habituate or adjust their responses over time, making it difficult to determine whether observed behavioural consistency reflects stable personality traits or simply learned test responses. Ensuring that these behavioural tests are both reliable and valid is therefore critical for accurately identifying personality traits in dairy cows.

The overarching objective of this experimental study is to enhance the understanding of personality traits in dairy cows and the methods used to measure them through behavioural tests and observations. To achieve this, the study focuses on several key objectives: first, identifying the key personality traits of dairy cows by using behavioural tests such as the Human Approach Test (HAT), Sudden Test (SUD), Novel Object Test (NOT), Open Field Test (OFT), and Novel Surface Test (NST). Second, establishing the validity of the set of behavioural tests used in this study by conducting repeated assessments to validate their short- and mid-term consistency, thereby evaluating the stability of personality traits. Third, evaluating the effectiveness of these tests in categorizing dairy cow personalities to deepen the understanding of herd social dynamics. We hypothesize that behavioural variables from different tests, shown to assess the same personality trait in dairy cows, will exhibit significant positive relationships, suggesting that these tests consistently capture similar personality dimensions. Additionally, we expect that cows’ reactions to environmental stimuli, such as their reactivity, exploration, and coping strategies in response to aversive situations, will remain relatively stable over time, reflecting the consistency of their underlying personality traits.

## 2. MATERIALS AND METHODS

### 2.1. Ethics Statement

The animals and research procedures used in this study were approved by the Animal Care Committee of McGill University and affiliated hospitals and research institutes (MCGL 10059).

### 2.2. Animals and Experimental Design

Lactating dairy cows from McGill University’s Macdonald Campus Farm (Sainte-Anne-de-Bellevue, QC) were selected for testing conducted in three periods from June 8^th^, 2023, to August 9^th^, 2023 (Figure 1). Of the total lactating cows housed at the farm in tie-stalls, a maximum number of individuals were included at each testing period. Participation varied due to differing parities (e.g., cows expected to become dry during an upcoming testing period were excluded) and physical conditions (i.e., sickness or injuries) across the three months (Period 1: 61 cows; Period 2: 69 cows; Period 3: 73 cows).

**Figure 1.**
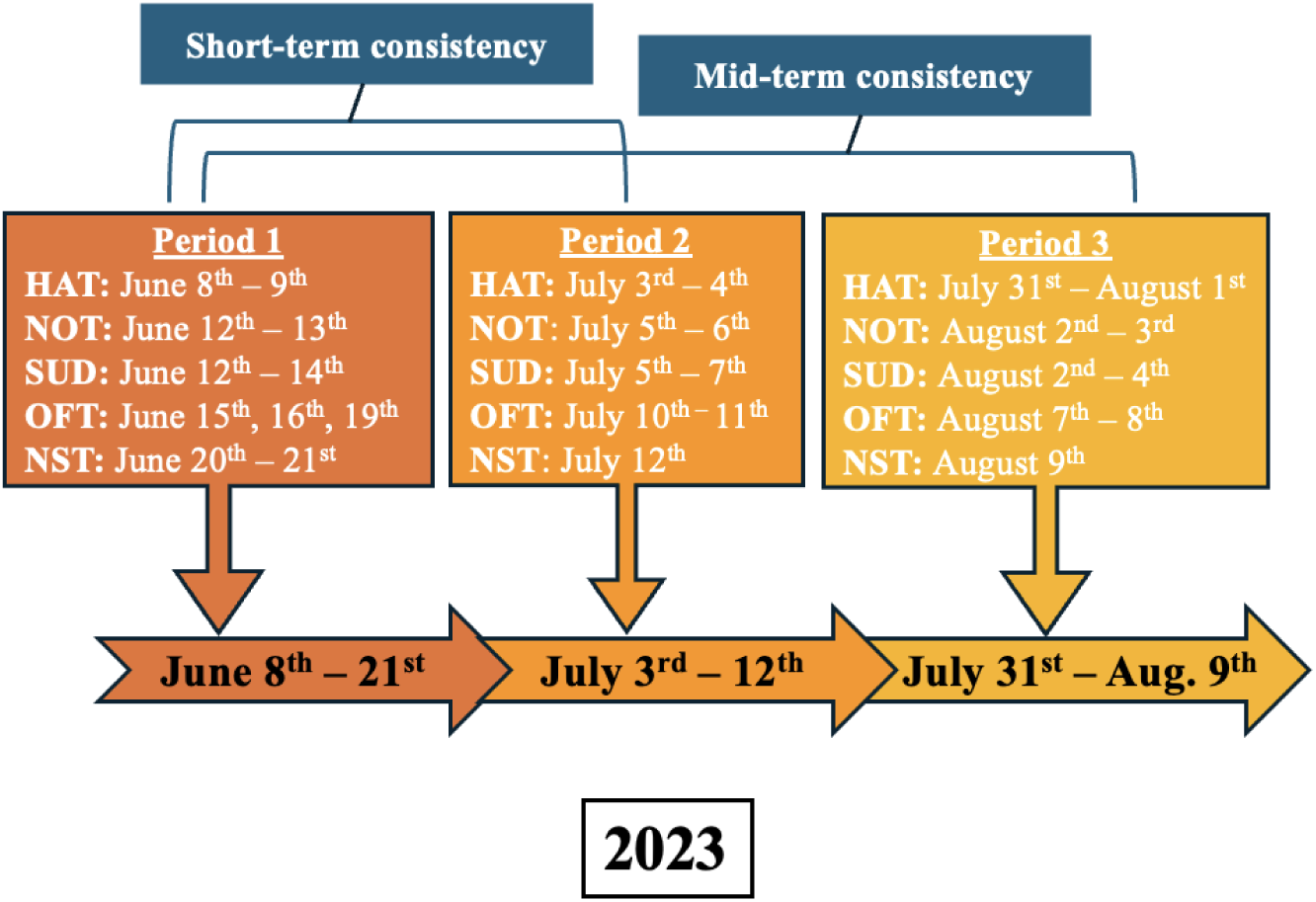
Testing was conducted between June and August 2023. The schedule also details the days allocated to each test. Acronyms for each test are as follow: Human Approach Test (HAT), Novel Object Test (NOT), Sudden Test (SUD), Open Field Test (OFT), Novel Surface Test (NST).

No treatments were applied to the cows before or during testing to ensure that their natural behaviours in response to stimuli remained unaffected by prior habituation or external influences. The cows were housed in individual tie-stalls measuring 186 cm by 142 cm, with tie rails positioned at a height of 122 cm and 36 cm from the manger wall. The flooring consisted of concrete rubber mats (KKM longline, Distribution Multi-Mat Inc., Ste-Cecile-de-Milton, QC, Canada), and approximately 2 cm of wood shavings were used as bedding, replaced as needed. Cows had daily access to an outdoor area, except during periods of extreme heat when outdoor access was restricted. Cows had ad libitum access to water and were fed a total mixed ration (TMR) four times daily. Milking occurred twice daily at 12-hour intervals (5:00 AM and 5:00 PM).

Three tests were conducted on all individuals during each testing period to assess personality: the Human Approach Test (HAT), the Sudden Test (SUD), and the Novel Object Test (NOT). These tests were conducted directly in the cows’ home stalls to assess responses in a familiar environment and avoid handling stress. Cameras were either mounted on tripods positioned in front of the stalls or attached to the barn walls, depending on the available space around each stall. A subsample of thirty cows, consistently available across all three testing periods (i.e., not dry, sick, or injured; one cow was removed in Period 3 as she had dried), were selected to participate in two additional tests: the Open Field Test (OFT) and the Novel Surface Test (NST), which aimed to create a more comprehensive personality profile. The OFT and the NST were performed outside the housing area: the OFT was conducted in an enclosed indoor arena within the barn, while the NST took place in a fenced outdoor corridor adjacent to the barn that served as the access route to the cows’ outdoor exercise area and was therefore familiar to the animals. This arrangement ensured that cows were exposed to both familiar and novel environments across the different tests. The order of tests remained consistent across periods (Figure 1), with the HAT always conducted on the first day to minimize potential carryover effects from prior interactions with researchers, ensuring cows were in a neutral routine state when encountering the unknown human. The NOT and the SUD were then conducted for all cows, but no cow underwent both tests on the same day. These tests were performed approximately one week before the OFT and NST, as they were logistically simpler and allowed for group testing. The OFT and NST were conducted on subsequent days. To minimize potential confounding effects between tests, each cow underwent only one test per day, reducing the risk of behavioural carryover that could bias observations.

All tests, except for the OFT, were recorded using GoPro® HERO7 Black and GoPro® HERO11 Black cameras (GoPro Inc., San Mateo, CA, USA) that were securely attached on either tripods or wall mounts. For the OFT, a set of seven high-performance cameras (Basler Ace, Ahrensburg, Germany) were positioned on the ceiling (2.3 m above the ground). Each camera was installed to ensure overlapping views with at least two other cameras, allowing all angles to be captured. All cameras were connected via wired connections to a desktop computer, and video capture was facilitated using the CONTEMPLAS TEMPLO capture engine (CONTEMPLAS GmbH, Kempten, Germany) to ensure synchronized recording of all videos.

The daily average temperature, particularly during periods of extreme heat, was documented as a variable potentially influencing cow behaviour. The average temperature during testing periods in degrees Celsius (C°) were as follows: Period 1: 21.4 ± 3.8; Period 2: 26.4 ± 2.6; Period 3: 21.2 ± 2.5. Wind velocity was measured specifically for the NST, the only test conducted outdoors, with average wind velocities (in mph) recorded as: Period 1: 5.5 ± 0.5; Period 2: 7 ± 0.5; Period 3: 5 ± 0.5. All weather data were sourced based on conditions in the Montreal area.

### 2.3. Human Approach Test (HAT)

The objective of the HAT was to study the animal’s response to an unfamiliar human approaching in a familiar environment (e.g. Aigueperse & Vasseur, 2021).

Throughout all three periods, an unknown human (UH), who had no prior interaction with the cows and had not visited the barn before, conducted the test, following the procedure described in Wolfe et al. (2026a). Each test was brief enough that the cows were not expected to become familiar with the UH during the session. The UH, a male standing at 175 cm tall, consistently wore white coveralls with the hood drawn up, white latex gloves, a light blue surgical face mask, and black rubber boots. The same human was used across all periods but did not participate in any other tests or interact with the cows outside of the testing context. This design aimed to reduce the likelihood that behavioural consistency (or lack thereof) was driven by familiarity with the stimulus itself. Using a single standardized human also limited additional variability in responses that could arise from differences in human appearance, posture, or behaviour. The observer, dressed in navy overalls and black rubber boots, remained the same individual for all sessions. Both the UH and observer underwent training by an experienced individual familiar with the procedure and practiced by observing cows not involved in data collection.

A testing sequence was established to avoid testing neighbouring cows within short intervals, which could potentially influence their behaviours. The following procedures were adapted from the methodology of Aigueperse & Vasseur (2021).

Prior to the start of testing, the test cows were prompted to stand up in the designated sequence by gently nudging them on the rump if they were lying down. Approximately 1 minute was allowed for cows to stand up and complete physiological processes such as the need to defecate or urinate, ensuring their focus remained on the test. Testing then proceeded according to the established order. The observer positioned themselves at least two stalls away from the focal test cow and ensured they did not disrupt other cows awaiting testing.

After the preparation phase, the UH adopted the starting position with arms at their sides and head facing the cow, positioned approximately 2 meters in front of the animal. The UH attracted the cow’s attention by using gentle clapping or snapping gestures if the cow was engaged in activities where her focus was not on the human, such as eating. Each test, lasting 25 seconds, consisted of 5 stages, with 5 seconds allocated to each stage, progressively involving closer or more direct interactions with the cow. The procedure was repeated two additional times per cow, totaling three tests per cow per period. Repetitions were conducted non-successively, such that a cow’s second and third tests occurred only after all cows in the testing order had completed the preceding repetition, with at least a 15-minute interval between repetitions. If at any stage the cow exhibited aggressive behaviour or posed a potential risk to the UH, the observer immediately declared “end test” and terminated the procedure.

### 2.4. Novel Object Test (NOT)

The objective of the NOT was to assess cows’ behavioural responses to novel objects within a familiar environment, and the test was conducted following the standard operating procedure described in Wolfe et al. (2026b).

Three objects with distinct shapes and colors but consistent textures (either made of ropes or uniformly wrapped in rope) were selected to ensure novelty while controlling for some aspects of palatability (e.g., preferences for rougher or smoother textures among cows) across all periods.

In Period 1, dog toys of various colors (green, orange, pink, and blue) consisting of tightly woven ropes with a spherical knot (diameter: 12 cm) and a loop handle (length: 19 cm) were used (see Figure 2A). For Period 2, PVC pipes (diameter: 18 cm, length: 45 cm) covered in diamond braid polypropylene ropes (3 colors: red, white, or blue; diameter: 9.5 mm) were employed (see Figure 2B). In Period 3, yoga blocks (length: 15 cm, width: 10 cm, height: 12 cm) wrapped in ropes of the same type and colors as those covering the PVC pipes were utilized (see Figure 2C).

**Figure 2.**
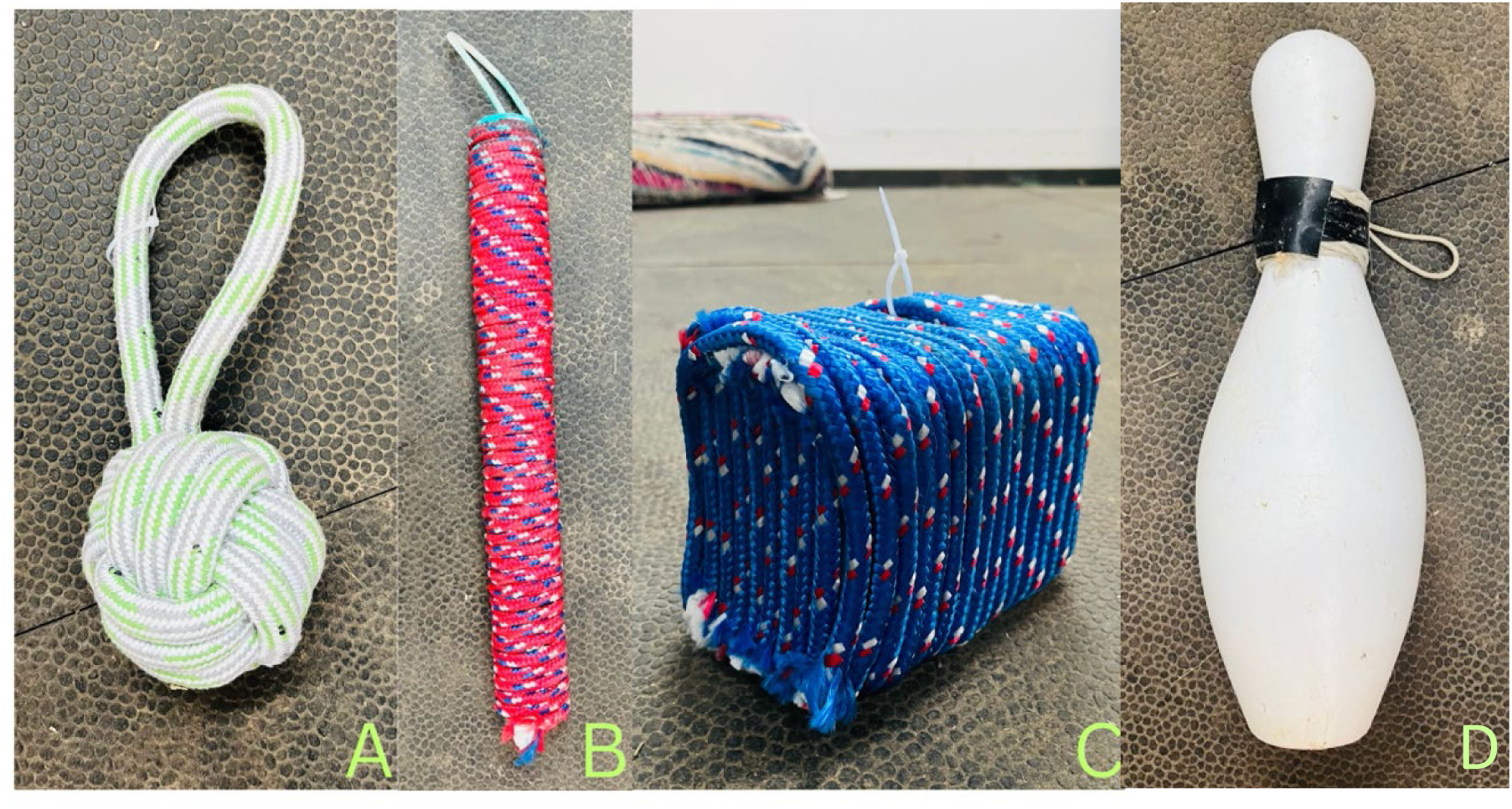
Novel objects used across testing periods. **(A)** Period 1: Rope-made dog toys. **(B)** Period 2: PVC pipes wrapped with polypropylene rope. **(C)** Period 3: Yoga block wrapped in polypropylene rope. **(D)** White hollow bowling pin used for the SUD tests.

The following testing procedures were adapted from the methodology of Herskin et al. (2004). To conduct the test, each test cow had an object clipped to the corresponding string in front of their stall. The object was then raised out of the cow’s view (Figure 3). This setup was completed in the morning before testing, during outing hours when the cows were in the exercise yards, to minimize disruption and reduce exposure to the object. Given the large number of participating cows, a streamlined approach was implemented for the NOT for more efficient management. Instead of lowering the object individually in front of each cow (as previously done by Herskin et al., 2004), the test was conducted row by row to optimize time, with objects lowered simultaneously in front of all cows within a row (n = 20 - 22 cows per row). Dividers installed during the preparation week minimized disturbance between neighbouring cows and prevented physical interaction with neighbouring objects, ensuring the test remained individual despite objects being lowered simultaneously.

**Figure 3.**
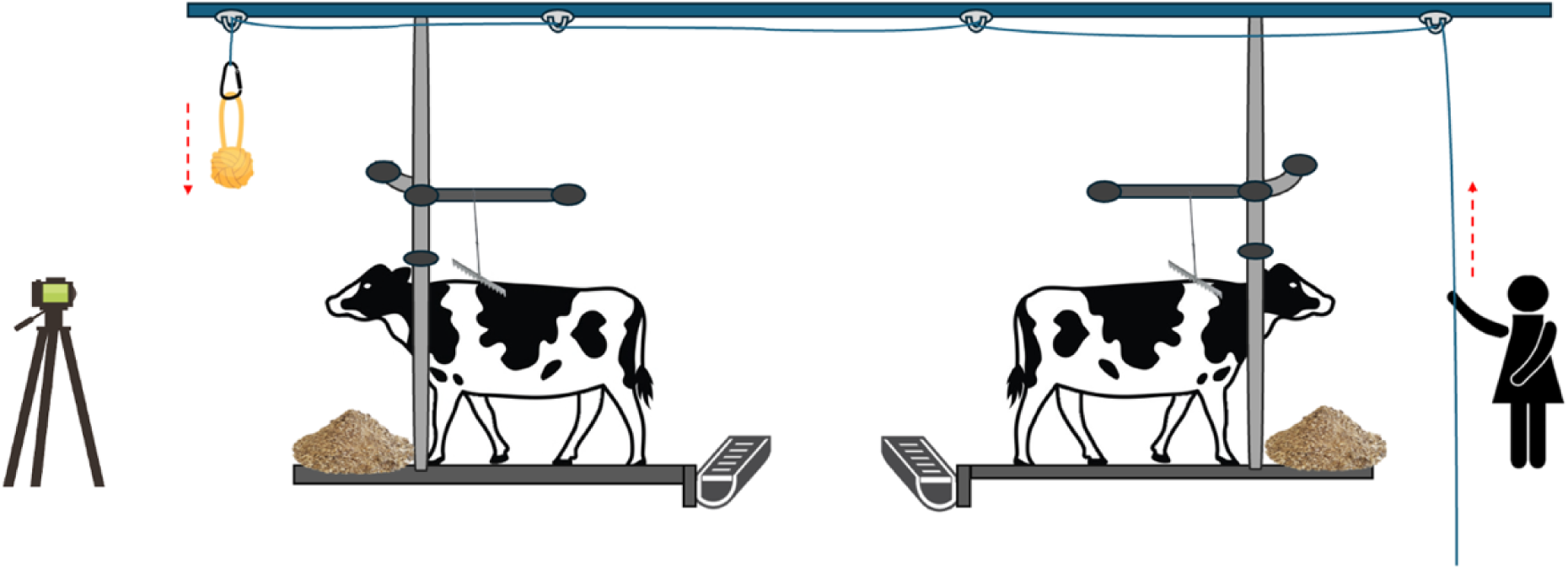
An illustration of the stalls from a side view featuring the string and pulley system. This method enabled each handler to simultaneously lower 2 or more objects, facilitating tests on 20-22 cows within a single blocked test period. To minimize human disturbance, objects were lowered overhead from the ceiling, while handlers remained out of sight behind the cows, separated by the middle aisle and a row of cows not being tested.

The experimenters ensured all test cows were awake (if not, they used their voice or gently tapped the cow to wake her up). A 2-minute waiting period allowed cows to settle, adjust, and complete physiological processes. If during the 2 minutes prep-time, a test cow went into a sleeping position (i.e., folding the head back against the flank) or appeared to be asleep, an experimenter would approach and gently tap her to wake her up.

After the 2-minute preparation time had elapsed, the experimenters lowered the novel objects in front of each cow at a gradual and uniform pace while striving to center the objects within each stall. Once all objects had reached the ground, cow behaviour was recorded for a 5-minute period.

### 2.5. Sudden Test (SUD)

The aim of the SUD was to assess the cow’s reactivity to a sudden event (e.g. Aigueperse & Vasseur, 2021). The test was performed using a white hollow plastic bowling pin (height = 45 cm, width = 10 cm, Figure 2D), dropped quickly in front of their stall (standard operating procedure described in Wolfe et al., (2026c)). For all periods, the same object was utilized due to its light weight and rounded shape, minimizing the risk of injury to the cows in the event of it falling on them.

Although the setup for the SUD was similar to that of the NOT (Figure 3), testing an entire row of cows simultaneously was avoided to prevent overreactions or responses influenced more by neighbouring cows rather than by the object itself. Instead, a chosen subset of cows (n= 1 – 5 cows), ensuring they were not adjacent, were tested simultaneously. In the end, each cow underwent a single SUD test per period, with no repeated testing within the same period; subsets were used only to structure test order. Following the testing of each subset, the next group was tested after an interval of approximately 5–10 minutes. The following procedure was adapted from the methodology of Aigueperse & Vasseur (2021).

Before the beginning of the test, the experimenters ensured that the focal cow and her neighbouring cows were standing to prevent potential injuries (such as focal cow stepping on a neighbour’s udder) should she react strongly to the sudden event.

Cows were required to remain standing for a total of 3.5 minutes: 2 minutes for preparation and habituation, followed by 1 minute and 30 seconds for the actual test before the object drop. Once this time elapsed, all experimenters released the ropes, quickly dropping the object in front of the cows’ stalls as centrally as possible. The object remained in position for 30 seconds before the test concluded.

### 2.6. Open Field Test (OFT)

The OFT was designed to evaluate an individual cow’s behavioural responses to an unfamiliar open environment and to social isolation. In some literature, the OFT is commonly conceptualized as a Novel Environment Test, as it exposes animals to an unfamiliar setting and assess their response to isolation, particularly in terms of exploration (Carter et al., 2012).

The Open Field Test arena consisted of a large enclosed fenced area (dimensions: 8.81 m × 6.55 m), with uniform maroon tarps secured from the ceiling to the fences. Flooring consisted of rubber matting and was consistent across all tests. A start box, defined by a triangular area enclosed by three fences (4.21 m, 4.10 m, and 2.71 m), was located at the entrance of the arena and separated from the main arena by a tarp doorway and removable drop bar, allowing controlled entry. The arena was otherwise empty and unfamiliar to the cows at the time of testing.

The test was conducted following Wolfe et al., (2026d) SOP which was adapted from the methodologies of Seaman et al. (2002) and Le Neindre (1989). Each cow was led by barn staff using a halter through the arena room door into the start box, where she remained for 30 seconds to habituate to her surroundings. After this period, the tarp doorway was rolled up and the drop bar was removed, allowing the cow to enter the arena. The cow had 30 seconds to enter the arena on her own; if not, encouragements, such as clapping and gentle pushing, were applied by one of the observers, who was kept consistent for all cows to avoid bias. If the cow entered the arena, an observer recorded her latency to exit the start box, defined as the time taken for both hind legs to fully cross the tarp curtain threshold. The drop bar and tarp door were then reset to their original positions to close off the entrance. The cow remained in the arena for 5 minutes, unless she escaped by knocking down the drop bar and re-entering the start box, in which case the test ended when she fully exited the arena. After the 5-minute testing period, the tarp door was raised, the start box swung open, and the drop bar was removed to allow the barn staff to enter the arena and lead the cow back to her respective stall. An illustration of the arena room setup can be found in Figure 4.

**Figure 4.**
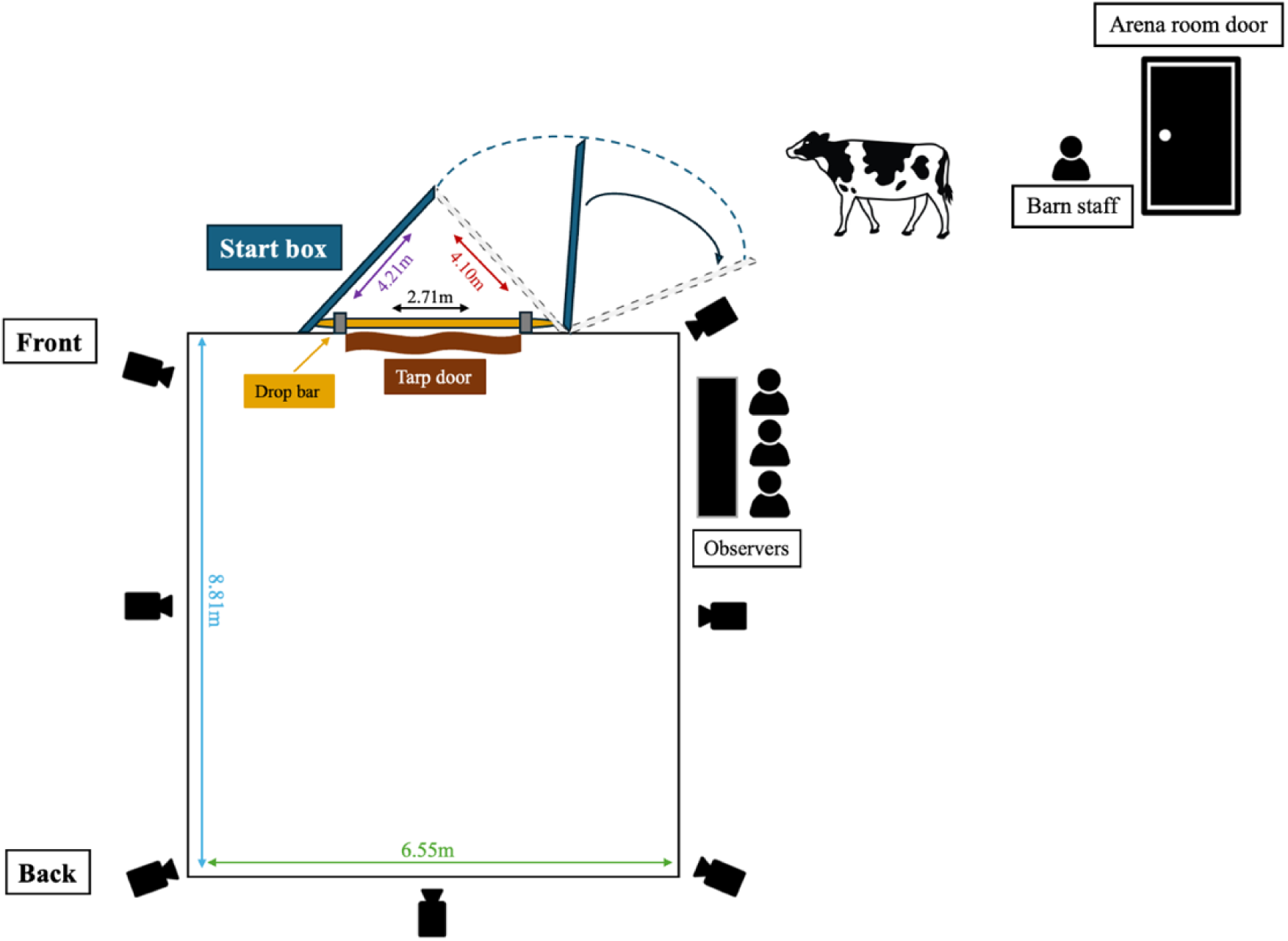
A diagram showing the testing arena room, the dimensions of the start box, as well as the arena itself.

### 2.7. Novel Surface Test (NST)

The Novel Surface Test (NST) exposed cows to an unfamiliar surface, eliciting behavioural responses to novelty similar to those assessed in the Novel Object Test (NOT). In addition to novelty, the NST required cows to cross the unfamiliar surface in order to move forward, thereby providing a context in which risk-taking behaviour, often discussed as a component of the bold– shy trait continuum, could be expressed. During testing, cows were removed from their stalls and placed individually in an outdoor corridor following the procedure described in Wolfe et al., (2026e); although the corridor itself was familiar, conspecifics were present beyond the novel surface. These animals, along with the outdoor area, served as contextual stimuli rather than experimental factors, and social motivation was not directly measured.

Two conspecifics were placed in the pen at the end of the corridor to utilize cow’s gregarious nature and encourage the test cow to cross the surface for social interaction.

Three surfaces with varying textures and appearances were selected for the three experimental periods. These surfaces consisted of everyday items that were non-slip and toxin-free, ensuring the safety of the cows while walking or exploring and minimizing the risk of injury.

Throughout all periods, the novel surface consisted of three wooden boards measuring 3.93 m in length and 2.24 m in width. It was positioned in Zone B of the corridor to cover the entire width of the testing corridor (approximately 3.23 m) (Figure 6). In Period 1, black grip tapes (width: 2 inches) were securely affixed to the wooden boards with a 2-inch gap between each tape, using wood glue (see Figure 5A). For Period 2, a conventional short-haired, multi-colored zebra-patterned carpet was cut and fitted onto the boards to achieve maximum coverage and was secured with nails (see Figure 5B). In Period 3, blue and purple yoga mats were cut into rectangular shapes (each measuring 61 cm in length and 30.5 cm in width) and arranged in a checkered pattern. These mats were nailed onto the boards (see Figure 5C).

**Figure 5.**
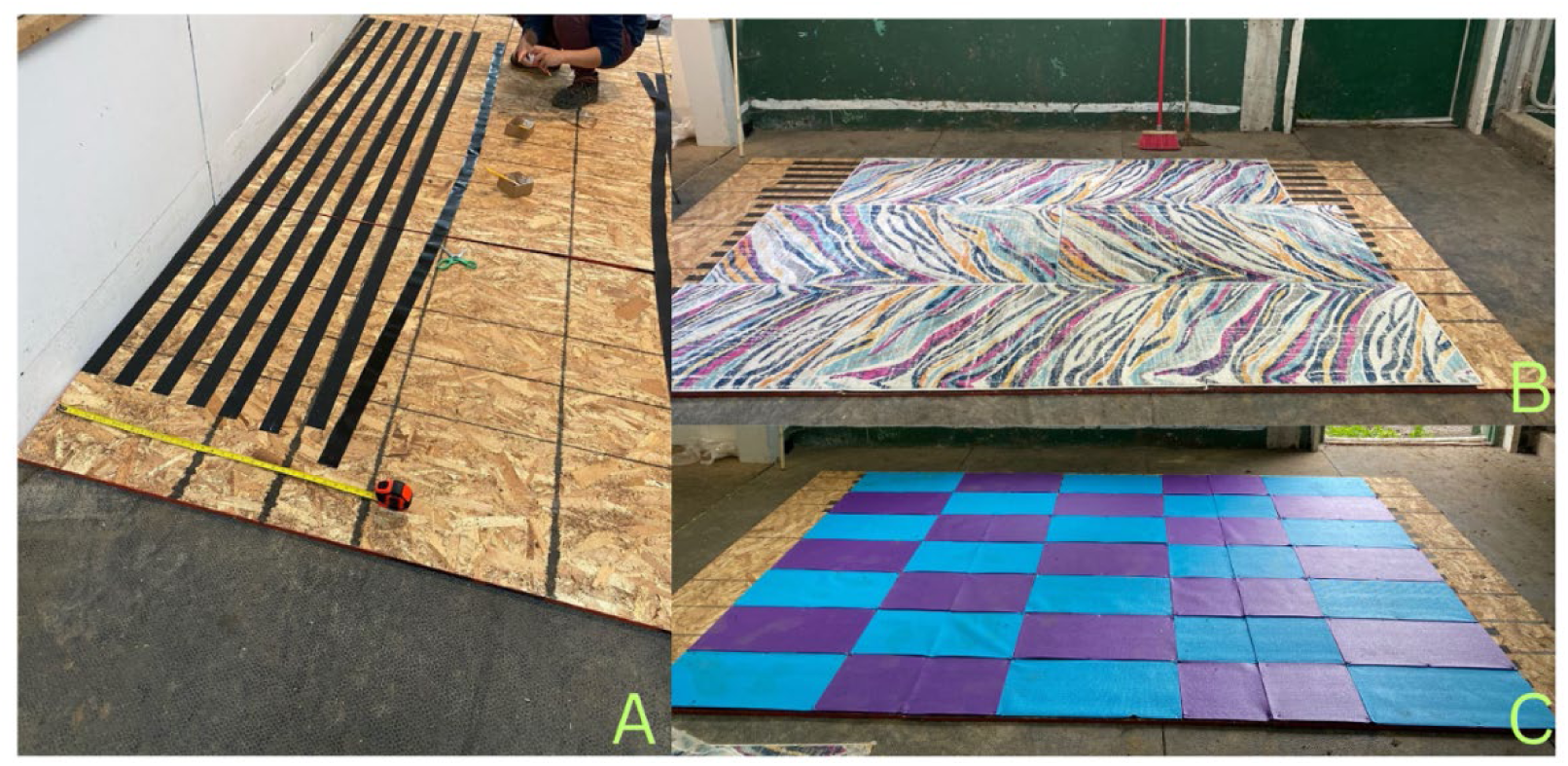
Novel surfaces constructed for each testing period. (A) Period 1: Wooden boards with black grip tape. (B) Period 2: Patterned carpets. (C) Period 3: Checkered blue and purple yoga mats.

**Figure 6.**
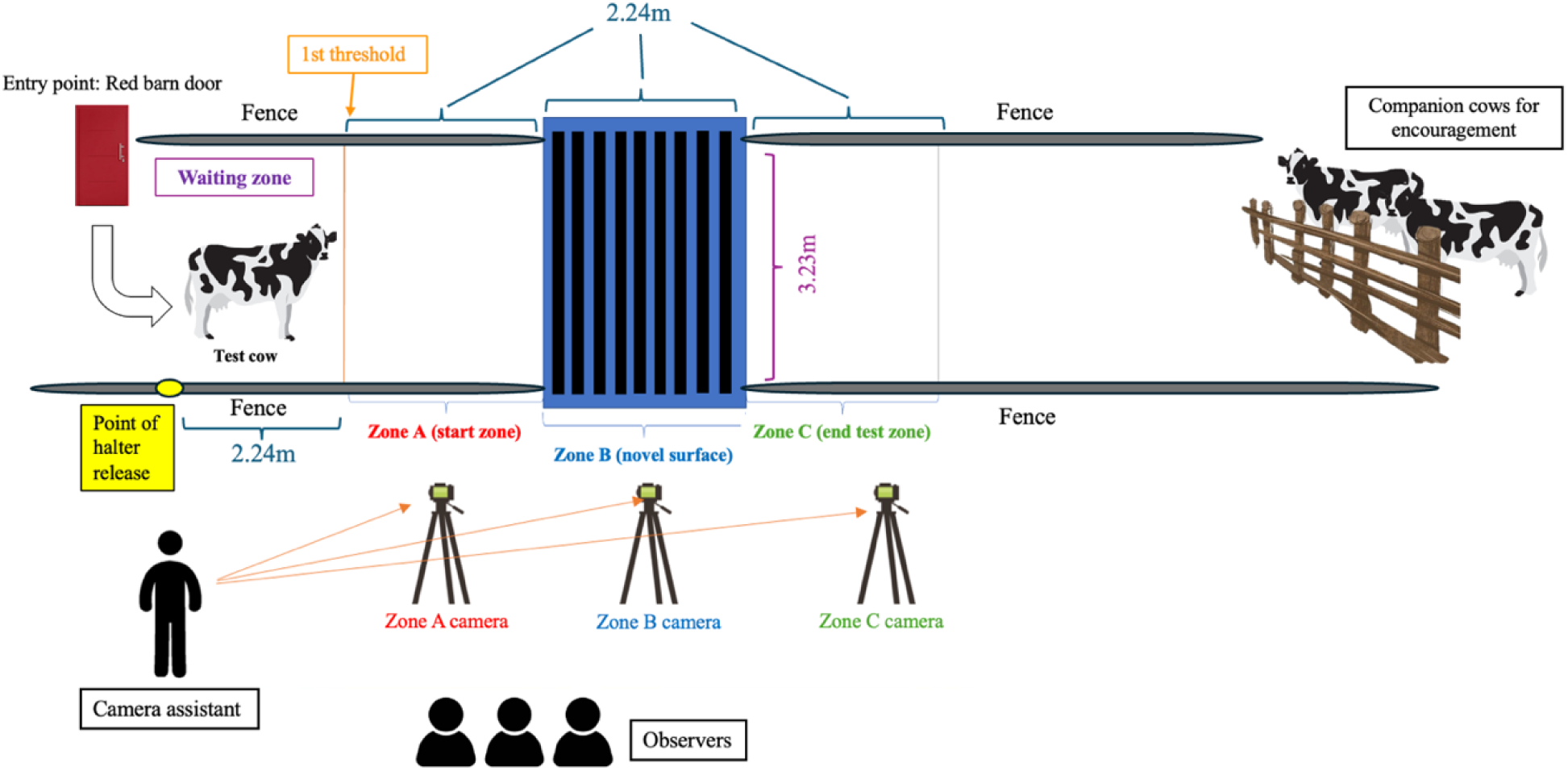
Illustration of the NST testing corridor layout, depicting zone locations, threshold markers, corridor dimensions, and the positioning of conspecifics. The waiting zone was subdivided into three equal sections.

Three GoPros mounted on tripods were used for recording (Figure 6): the camera in Zone A captured from the red barn door to the beginning of Zone B, the Zone B camera focused exclusively on the novel surface, and the Zone C camera recorded from the end of Zone B to the end of Zone C, including the paddock with the two companion cows to observe any social interactions with the test cow.

The following procedures were adapted from the methodologies of Lansade & Bouissou (2008) and Gonzalez-De Cara et al. (2017). Each cow was led through the barn door by a staff member using a halter. Upon reaching the designated point for halter release the barn staff released the halter and exited the corridor. There was a pre-test interval of up to 2 minutes from the barn staff’s exit until the cow crossed the first threshold, marked by a line of black tape, into Zone A. The test officially began once she crossed into Zone A.

If the cow did not cross into Zone A within the initial 2-minute interval, the barn staff re-entered the corridor and gently encouraged her forward using different non-aversive methods (e.g. soft clapping, light guiding pressure, or leading with the halter) depending on her responsiveness, to avoid eliciting a human-reactivity response unrelated to the test. If, after 2 minutes of encouragement, the cow remained in the waiting zone, the test would be stopped and deemed “unsuccessful”. The cow would then be led back to her stall.

Once the test began, the cow then had 3 minutes to cross the surface and proceed into Zone C. Following completion of the test, the cow was allowed a 20-second reward period to interact with her companions or explore the environment. The barn staff then guided the cow back to her stall.

### 2.8. General Behavioural Measures and Observations

#### 2.8.1. HAT Behavioural Analysis

For the analysis of the HAT, all behaviours and scores were recorded live by a trained observer following the scoring table outlined in Table 1 (adapted from Aigueperse & Vasseur (2021)). Prior to data collection, the observer was trained by an experienced observer using the same scoring protocol, and initial observations were conducted jointly to ensure consistent interpretation of behavioural categories.

**Table 1.** Scoring table used for the Human Approach Test.

| **-3** | **-2** | **-1** | **0** | **1** | **2** | **3** |
| --- | --- | --- | --- | --- | --- | --- |
| Steps back more than 2 steps, chain stretched to max (and/or hidden head) | Steps back (1 or 2 steps) / struggles (for the last stage only) | Turns head back or away | Looks at the person | Approaches the person without touching, sniffs | Approaches and touches the person | Tries to lick/catch the person with their mouth (clothing or hand with the tongue or the lips). |

For each stage, the observer assigned a score ranging from −3 to 3 based on the cow’s behaviour and noted the latency to voluntarily make contact with the unknown human, if applicable. In cases where the test was terminated early due to aggressive behaviour, cows did not receive behavioural scores for the remaining stages. These observations were treated as missing data, and “N/A” was recorded in all corresponding data cells. All results were subsequently combined and averaged to calculate a mean reaction score per stage across the three tests within each period, which was then used to assess intra-test reliability. These scores were later categorized and ranked, as explained below in the *3. Statistical Analysis* section, to evaluate test validity, where greater deviations between periods could suggest a lack of consistency over time.

#### 2.8.2. SUD Behavioural Analysis

For the analysis of the SUD, cow reactivity was analyzed using video recordings, which were scored by a single observer. Prior to scoring, the observer was trained using previously recorded SUD videos and a gold standard video to ensure accurate and consistent application of the scoring criteria. Intra-observer reliability was subsequently assessed by rescoring a subset of videos, yielding a Kappa value of 0.90, indicating high consistency. Detailed scoring criteria are provided in Table 2 (adapted from Aigueperse & Vasseur (2021)). As with the HAT, these scores were subsequently categorized and ranked to evaluate temporal consistency.

**Table 2.** Reaction scoring table used for the Sudden Test.

| **0** | **1** | **2** | **3** | **4** |
| --- | --- | --- | --- | --- |
| No reaction | Startled, with no backward movement | Startled, with backward movement of 1 or 2 steps | Startled, with strong backward movement of more than 2 steps and/or taut chain | Startled, with strong backward movement of more than 2 steps and a taut chain, and either struggling OR moving diagonally backwards (AND/OR head is hidden) |

#### 2.8.3. NOT, OFT, and NST Behavioural Analysis

For the analysis of the NOT, OFT, and NST, the Observer XT 14 software (Noldus, 1991) was utilized to annotate the movements and behaviours exhibited by test cows from their recorded videos. All observers (n = 7 for NOT, n = 6 for OFT and NST) underwent standardized training using both a training video and a gold standard video. Annotation began only after each observer reached intra- and inter-observer reliability scores of 90% or higher. For each test, the duration (d), frequency (frq), and latency (lat) of all relevant behaviours were recorded (unless otherwise stated in the corresponding ethograms). Live observations recorded during the OFT (e.g., the latency to exit the start box, the frequency and pitch of vocalizations) and the NST (e.g., social interactions with companion cows, vocalizations, and test duration) were manually logged into Excel for future reference. Specifically for the OFT, the positions of the cow within the arena were scan-sampled at timed intervals. Frequency, percentage, duration, and latency measures were extracted for all behaviours; however, not all measures were included in the statistical analyses. Although a comprehensive set of behaviours was initially recorded, some variables were excluded prior to statistical analysis if they were rare, invariant across individuals, or unsuitable for multivariate analysis. Inclusion criteria and variable selection are described in *Section 3. Statistical Analysis*. Full ethograms and behaviour definitions are provided in Tables 3 (NOT), 4 (OFT), and 5 (NST).

**Table 3.** Ethogram of behaviours observed during the NOT (adapted from Herskin et al., 2004).

| Observation Category | Behaviour | Description |
| --- | --- | --- |
| **Behaviours related to novel object (d, frq, lat)** | **Approach** | The cow decreases the distance between herself and the object by moving her head towards the object. |
|  | **Move away/standing up** | Cow increases the distance between herself and the object by diverting her head quickly away (and/or stepping back or away and/or standing up from a lying position (i.e., where the cow is not bearing weight on her hooves and the majority of her body is making contact with the base of the stall, then lunges forward to displace her weight while lifting her back legs to rise and make contact with her hooves to the stall)). |
|  | **Sniff** | Muzzle close (less than 1 head of distance away) or in contact with the object with contraction of the nostrils. |
|  | **Lick** | Cow makes contact with the object using her tongue. |
|  | **Push/bite** | Head, including the muzzle, is in contact with the object to push or roll the object “intentionally”, or teeth is in contact with the object. |
| **Behaviours not directly related to novel object/normal behaviours (d, frq, lat)** | **Exploration** | The cow is chewing, licking, and/or sniffing non-feed objects (such as rope, dividers, chains, poles, etc., excluding novel objects). If nostril movement is not visible, the cow is considered to be exploring if her muzzle is close to a non-feed object for at least 3 seconds and she is not performing another behaviour. |
|  | **Sleeping** | Cow has eyes closed, with or without being in a lying position, and remains inactive for at least 5 seconds (head can be folded against body or not). No rumination visible. |
|  | **Eating/Drinking** | Searching, sniffing, grabbing, chewing and/or swallowing feed, or when the muzzle is located in the water bowl, with ingestion of fluids (including nostril licking after consumption). |
|  | **Social Interaction** | Cow makes intentional physical contact with another cow in the neighbouring stall (e.g., allo-grooming using the tongue, fighting by head butting) or is sniffing the cow in neighbouring stall. |
|  | **Resting/Inactive** | No activity for at least 5 seconds with eyes open (including when cow is vigilant), can be ruminating and can be standing or lying. |
|  | **Standing up** | The cow lunges forward and lifts her hindquarters, shifting her weight to lift her back legs until her hooves make contact with the base/floor of the stall. She then lifts her front legs into contact with the base, so all four hooves bear her weight. This behaviour begins as the cow lunges forward. |
|  | **Lying down** | The cow first bends her forelegs so that her knees are in contact with the base of the stall. She then bends her hindlegs to lower herself from a standing position, where all four hooves are bearing weight and touching the base of the stall, to a position where none of her hooves |
|  |  | bear weight and her sternum or side makes contact with the base. This behaviour begins when both of the cow's knees are touching the base of the stall. |
|  | **Scratching/**<br>**Grooming** | Cow rubs head or body against part of the stall to scratch herself or uses tongue to lick her own fur. |
|  | **Rumination** | The cow is eating partially digested feed and is either regurgitating or chewing, without previously eating. |
| **Distances**<br>**(d, frq, lat)** | **< 1 head** | A distance of less than 1 head between the cow and the object, with her muzzle close to or touching the object and her attention focused on it. This behaviour should be coded when the cow is performing actions such as sniffing, licking, or pushing, indicating interaction with the object. |
|  | **1 to 2 heads** | When the cow is past the point of the cement barrier at the base of the stall, but where the cow's muzzle is not in contact with the object or not intentionally near the object to interact. |
|  | **> 2 heads** | When the cow is in line with or behind the cement barrier at the base of the stall, but the chain is not stretched to the max. |
|  | **Stretched maximum** | The cow is at the greatest distance from the object, with the chain stretched to the fullest extent. |
| **Other behaviours**<br>**(d, frq, lat)** | **Looking at object** | When the muzzle is directed towards the object (if eyes aren't visible). If the whites of the eyes are visible, then it's when either the eyes, the muzzle, or both, are directed towards the object. |
|  | **Not looking at object** | Eyes and muzzle are not directed toward the object. |

**Note.** *d* = duration; *frq* = frequency; *lat* = latency.

**Table 4.** Ethogram of behaviours observed during the OFT, adapted from Seaman et al. (2002) and Le Neindre (1989).

| **Category** | **Behaviour** | **Description** |
| --- | --- | --- |
| **Movement (d, frq, lat)** | **Walking** | Cow moves forward at a slow and steady pace (for at least 3 steps or more). |
|  | **Trotting** | Cow moves at a quick and steady pace (for at least 3 steps or more). |
|  | **Vigilance** | Cow stops all movement and appears tense in the body and ears are facing forward, with eyes wide open staring at a point of interest. |
|  | **Charging** | The cow quickly approaches and aggressively pushes the tarp doorway and/or the pole as a mean to exit. |
|  | **Stationary (Body)** | Cow remains still for at least 3 seconds with a relaxed body posture and can be looking around and/or exploring. |
|  | **Pacing** | Repetitive locomotion back and forth along a wall repeated for at least twice. May be interspersed with escape attempts. |
|  | **Jumping** | Cow launches herself forward onto her two front legs (which can occur simultaneously or at different times). This behaviour ends when the cow's two front legs have landed on the floor and the body is well balanced and stable (there may be a sequence of jumps or one jump). |
| **Behaviours (d, frq, lat)** | **Exploration** | Cow is investigating with head less than 1 muzzle away from the floor/doorway/walls. |
|  |  | Cow interacts with objects or surfaces other than the walls, the doorway, and the floor. |
|  | **Grooming** | The cow uses her tongue to lick any part of her fur, or she uses her head, hoof, or surrounding environment to rub against any part of her body. |
|  | **Observing** | The cow's head can be moving, body is not tense, and she is looking at something in her environment. |
|  | **Head poking** | The cow either pushes her head against or places her head through the doorway or makes contact with the drop bar with her head. |
|  | **Vocalization** | The cow stretches her head forward or upwards with mouth open to produce a call/sound. |
|  | **Defecation (frequency only)** | Cow lifts tailhead before eliminating feces from her body. The behaviour ends once the waste has been discharged and the tailhead returns to a relaxed position. |
|  | **Urination (frequency only)** | Cow arches back and lifts tail to expel urine from her body. This behaviour ends once all the urine has been discharged and the tail returns a relaxed position. |
| **Rumination (d, frq, lat)** |  | Regurgitation, chewing and/or swallowing of previously eaten feed. |
| **Zones (d, frq, lat)** | **Door**<br>**Wall**<br>**Middle** | To be annotated by scan sampling every 30 seconds, initial scan starts at 0 second.<br>**Door/Wall/Middle:** Consider the cow in a zone when her 2 front legs are within a certain zone and it's less than a 1 body width away from the borders (if in doubt, e.g., when the 2 front legs are in 2 different zones, check the head position of the cow). |

**Note.** *d* = duration; *frq* = frequency; *lat* = latency.

**Table 5.**
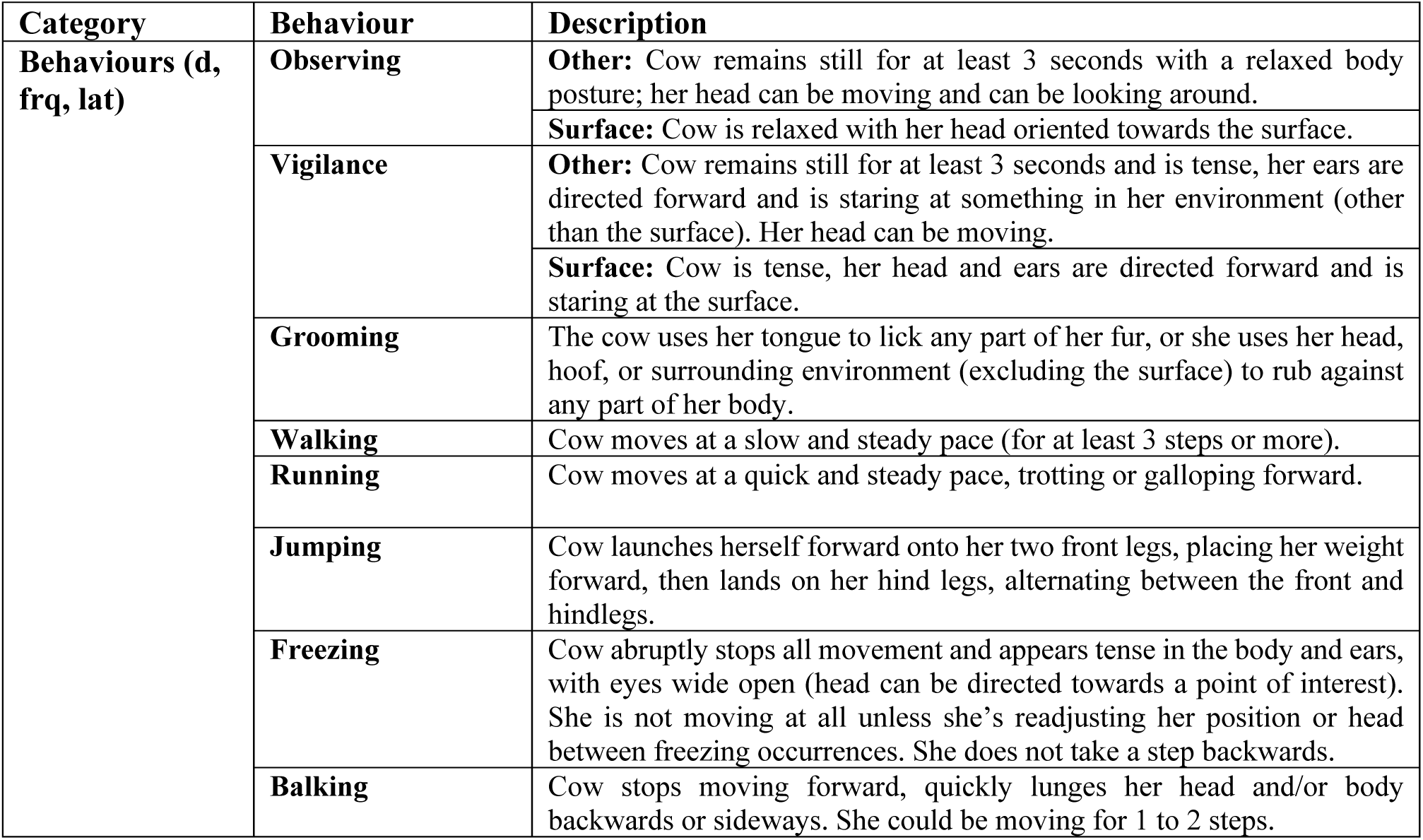
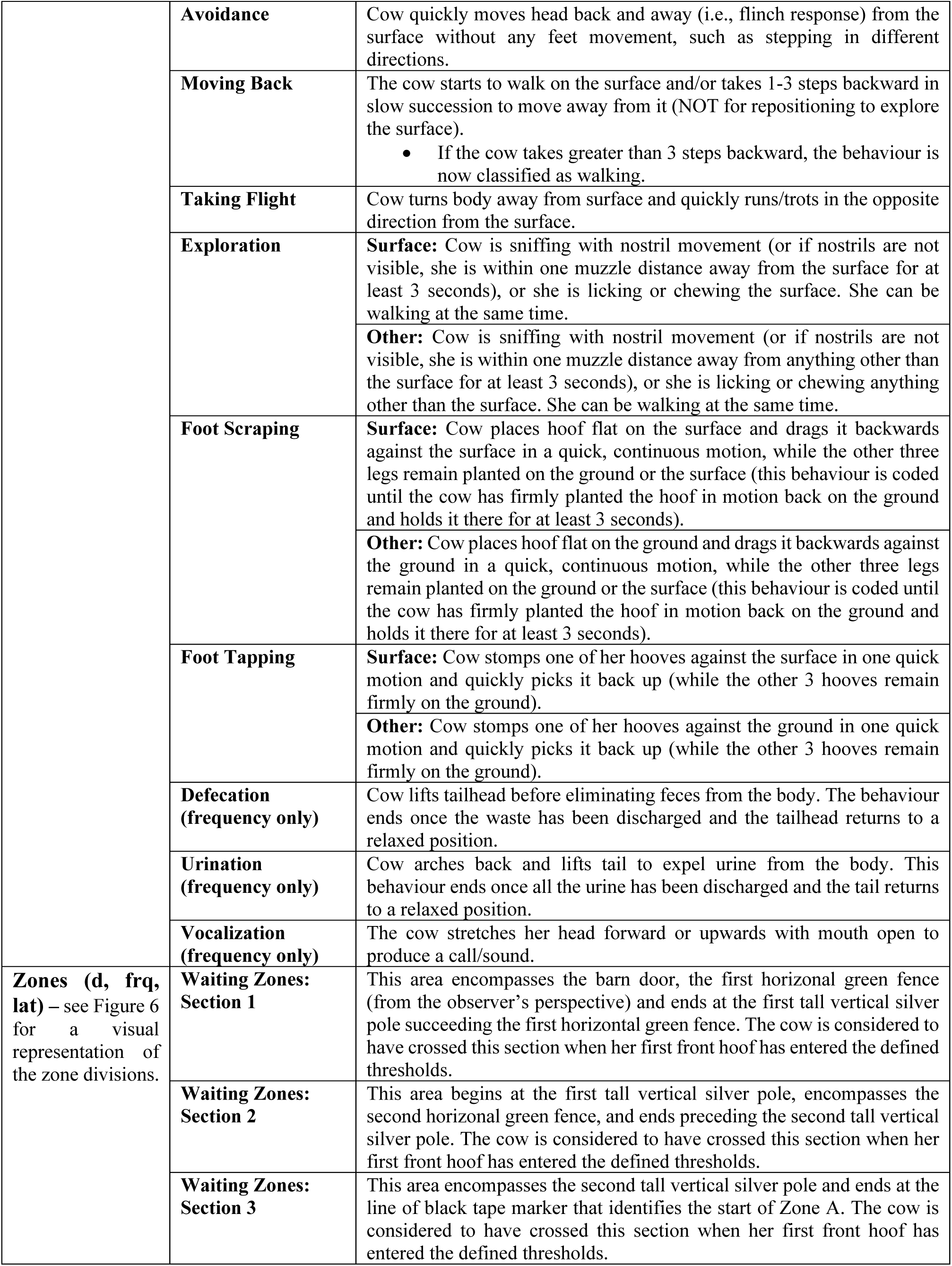
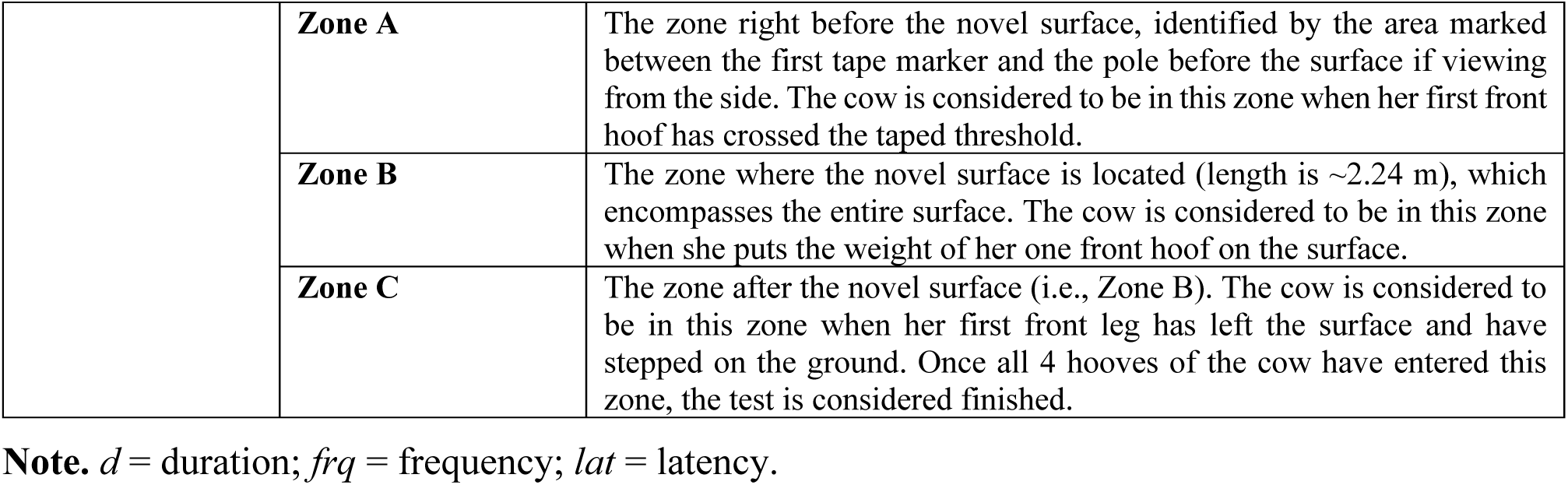
Ethogram of behaviours observed during the Novel Surface Test (adapted from Lansade & Bouissou, 2008; Gonzalez-De Cara et al., 2017).

## 3. STATISTICAL ANALYSIS

### 3.1. General Data Handling and Manipulation

Statistical analysis was performed using R (version 4.3.3; R Core Team, 2023) in R Studio (version 2023.12.1.402). The following packages were utilized (excluding base R packages): *dplyr* (for data manipulation, including filtering, selecting, and mutating), *car* (for data visualization through QQ-plots), *ggplot2* (for advanced data visualization), and *gridExtra* (for organizing and displaying multiple plots). Packages specific to individual tests and analyses are detailed in the sections below.

### 3.2. Data Manipulation for the Human Approach Test (HAT) and Sudden Test (SUD) by Categorization

A categorization method was used to organize reaction scores from both the HAT and SUD. This strategy facilitated the identification of patterns and trends, offering an advantage over the use of raw scores. As ties between individuals are common due to the ordinal nature of scores, categorizations reduce the impact of small fluctuations that might otherwise cause large rank shifts, ensuring that ties have less influence on the correlation.

By categorizing the data in the same way for both the HAT and the SUD, direct comparisons between the two tests become easier to interpret. This approach facilitates the identification of general behavioural trends, such as whether cows that display boldness in the HAT also exhibit similar traits in the SUD. In doing so, it provides valuable insight into patterns of personality consistency across tests and supports the evaluation of inter-test reliability.

In the analysis of the HAT, the original scoring system (Table 1) was reorganized into three categories reflecting levels of behavioural responsiveness to the human. This reorganization emphasized the magnitude of the behavioural response rather than its emotional valence. While the original scoring system distinguished between positive (approach) and negative (avoidance) reactions, these distinctions were intentionally collapsed to capture overall behavioural engagement with the human stimulus. The term ‘responsiveness’ was therefore adopted to describe the degree to which cows behaviourally reacted to the presence of an unfamiliar human, independent of whether the response was approach- or avoidance-oriented. This distinction was made to avoid conflating behavioural engagement with emotional valence. Conceptually, emotional reactivity broadly refers to how readily an individual experiences an emotion, as well as the intensity and duration of that response (Davidson, 1998; Zelkowitz & Cole, 2016), and is not limited to fear-related reactions (Mason, 1984; Boissy, 1995). This conceptualization aligns with broader frameworks emphasizing individual differences in behavioural coping styles and responses to environmental stimuli (Koolhaas et al., 1999):

**Category 1:** Score of 0 indicates no response towards the human.

**Category 2:** Scores ranging between −1 and 1 (excluding 0) represent minimal responses, with cows being responsive but not overly receptive towards human.

**Category 3:** Scores ranging from −3 to −1 (excluding −1) and from 1 to 3 (excluding 1) reflect moderate to strong response, encompassing behaviours such as cautious to pronounced avoidance, deliberate approach to intense engagement.

**Category 3** was created by merging two categories: Originally, Category 3 included scores ranging from −2 to −1 (excluding −1) and from 1 to 2 (excluding 1), while **Category 4** encompassed scores from −3 to −2 (excluding −2) and from 2 to 3 (excluding 2). Category 4 represented strong reactivity, either through extreme avoidance or intense engagement. However, because only a few individuals displayed these extreme behaviours, Category 4 was combined with Category 3. This adjustment was necessary due to the small sample size in Category 4.

For the SUD, the five reaction scores (shown in Table 2) were also grouped into categories based on the intensity of the cow’s reactions to the sudden drop of the object. Unlike the HAT, the SUD is designed to induce fear by presenting a sudden visual and/or auditory stimulus, which elicits both behavioural and physiological responses. As only the instantaneous reaction to the stimulus is annotated, the term “reactivity” would be appropriate to describe the cows’ behaviours in this context.

**Category 1**: Score of 0 indicates no reaction from the cow.

**Category 2**: Scores of 1 or 2 represent a moderate response, where the cow showed some reactivity to the sudden object but not an intense reaction.

**Category 3**: Scores of 3 or 4 indicate a heightened response, with the cow being startled and displaying a more intense behavioural reaction to the sudden object.

### 3.3. Data manipulation for the Novel Object Test (NOT), Open Field Test (OFT), and Novel Surface Test (NST) using Principal Component Analysis (PCA)

Principal Component Analyses (PCAs) were performed separately for each behavioural test (NOT, OFT, and NST) to identify patterns and reduce data complexity, aiming to explore relationships between the behavioural variables recorded for each cow within each test. Behavioural data, annotated using video analysis software, were expressed as frequencies per second, percentage of time, or latencies, depending on the nature of the behaviour observed. The general procedure for conducting PCAs is described below.

#### 3.3.1. Raw Data Manipulation and Adjustments Prior to Creation of PCAs

As a preliminary step, raw data from each test were refined by excluding variables lacking biological relevance or significance. For instance, behaviours like “sleeping” or “urination” were omitted, as they were either biologically meaningless in the context or too rare to influence PCA components. Additionally, mutually exclusive variables (e.g., rumination present and rumination absent) were evaluated for removal, as the presence of one inherently excludes the other. The retained variable was chosen based on its biological relevance to defining meaningful dimensions in the PCA.

After cleaning the raw variables, time associated with behaviours excluded from analysis was retained as unscored time rather than analysed as separate behavioural categories. For duration-based measures, the durations of all included behaviours were summed, and the proportion of time allocated to each behaviour was calculated relative to this total. This approach ensured that percentage values reflected the distribution of observed behaviours without distortion from unscored periods (e.g., technical interruptions or test termination conditions), thereby preventing misinterpretation of behavioural time budgets.

Due to technical issues during testing and some cows ending the tests prematurely, frequency-based variables were standardized by dividing raw occurrence counts by the total test duration, yielding rates per second. Duration-based variables were expressed relative to test duration (as percentages). This modification ensured that both frequency and duration measures were comparable across cows, accounting for variation in test durations.

In addition to standardizing and cleaning raw behavioural variables, certain derived measures were calculated from the processed data to better characterize aspects of cow movement relevant to subsequent PCA analyses. To characterize cow movement in the OFT, scan-sampled position data were used to derive a measure of zone transitions. A transition was recorded each time a cow moved to a different zone between two consecutive scans. This measure served as a representation of activity within the arena, reflecting changes in spatial location over time.

#### 3.3.2. Review and Refine Data Frame for Biologically Relevant Behaviours

Behaviours that do not offer substantial contributions (i.e., rare behaviours) may be considered for removal in this process. To further refine the dataset, a data frame was generated to pair variables with high correlations (|r| > 0.8). Decisions regarding the retention or removal of variables are guided by their biological significance. Highly correlated variables, which often introduce redundancy by providing overlapping information, are carefully assessed. Both variables may be retained if they offer valuable insights, with final decisions postponed until PCA visualization clarifies their contributions. This approach ensures a balance between reducing redundancy and preserving essential information.

Principal Component Analysis (PCA) is a type of Biplot Component Plot that simplifies complex datasets by identifying patterns in variable correlations and reducing dimensionality. By transforming high-dimensional data into principal components, PCA captures essential trends, minimizes redundancy, and reduces noise, leading to clearer and more interpretable results. To determine if a dataset is suitable for dimensionality reduction, the Kaiser-Meyer-Olkin (KMO) test is conducted using the *corpor* package in R Studio, which regularizes the correlation matrix and performs the KMO assessment. This test evaluates the adequacy of the sample by measuring the shared variance among variables. The KMO statistic, ranging from 0 to 1, indicates the dataset’s appropriateness for PCA, with values below 0.5 signaling weak correlations that may hinder its effectiveness. While the KMO test ensures statistical viability, the biological relevance of the variables should still be considered when making decisions about their inclusion.

#### 3.3.3. Creating PCAs for Visualization and Naming Dimensions

Using R Studio and the *Factoshiny* package, a PCA graph representing all behavioural variables (excluding those previously removed) was generated. If necessary, behaviours were grouped based on their loadings and positions within specific quadrants, aligning variables with conceptual or functional similarities and biological relevance. The grouping of behaviours was not performed arbitrarily, but followed a two-step process combining statistical structure and biological interpretation. First, all raw variables were included in an initial PCA to identify behaviours that clustered together based on shared variance. Behaviours loading closely within the same region of the PCA space were then evaluated for conceptual similarity, and when both statistical association and biological meaning aligned, they were combined into a composite variable. To maintain methodological consistency, only behaviours represented using the same unit (e.g., percentage of time) were grouped together. When a grouped variable was retained in the final PCA, the original individual variables used to construct it were excluded to avoid redundancy and double representation of the same behavioural information.

Based on this approach, the grouped behaviours for each test are as follows: In the NOT, object-oriented behaviours (represented as *P_ObjectOriented* on the biplot) include licking, sniffing, biting, and pushing the object. These behaviours were grouped to capture overall engagement with the novel object, reflecting the cow’s motivation to interact with the stimulus rather than avoid it. Although these behaviours differ in their intensity and proximity of interaction (e.g., cautious investigation versus close physical contact), they were retained within a single category because they loaded closely together in the PCA space, indicating strong correlation and shared variance. This grouping suggests that, within the context of the NOT, these behaviours collectively represent a common behavioural tendency toward object-directed engagement. Avoidance behaviours, labeled *P_Avoidance*, include more than 2 heads away from the object, chain stretched to the max, and move away/stand up. In the OFT, fear-related behaviours (*Freq_Fear-related Behaviours*) such as vigilance, vocalization, and jumping are grouped together to reflect behavioural responses associated with heightened arousal in the presence of a potentially stressful environment. High arousal locomotory behaviours (*Freq_Locomotory High Arousal Behaviours*) such as charging, trotting, and pacing were grouped to capture intense movement patterns indicative of elevated activity levels. Although some behaviours (e.g., jumping) may possibly reflect both fear-related responses and high-arousal locomotion, behaviours were categorized based on their predominant functional context within the OFT. These behaviours were retained as separate variables prior to analysis, and their categorization was supported by their close association and shared loadings within the PCA space, indicating that they co-varied as part of distinct behavioural response patterns. For the NST, behaviours related to engagement with the surface, such as Exploring the surface and Foot Tapping, are grouped under *Perc_Engagement Surface*. Both behaviours were quantified as duration-based measures reflecting the time cows spent actively interacting with the surface. These behaviours were not combined into a single metric; rather, they were retained as separate variables and interpreted together based on their shared loadings within the PCA, indicating a common tendency toward surface-directed engagement. Lastly, time spent in the waiting area (*Perc_WA_TOT*) includes being in waiting area sections 1, 2, and 3. Latency measures were not retained for PCA analyses. Although latency to first contact or approach is commonly reported in novelty tests, latency variables are often truncated by the test duration when animals do not perform the behaviour within the test duration. As a result, latency showed limited variance and weak associations with other behavioural variables in preliminary exploration, reducing its suitability for multivariate analyses aimed at identifying patterns of co-occurring behaviours. The present analyses therefore focused on frequency- and duration (percentage)-based measures that better captured sustained behavioural responses throughout the test.

Additionally, when frequency- and duration-based measures described the same behavioural expression and were strongly correlated, only one metric (either frequency per second or percentage of time) was retained to avoid overrepresentation of the same behavioural tendency in subsequent analyses. This decision was made on a case-by-case basis, balancing statistical redundancy with biological interpretability.

Eigenvalues were extracted following the Kaiser criterion, which recommends retaining components with eigenvalues greater than one. This criterion was used as an initial guideline for dimension retention. When more than two components met the eigenvalue threshold, the number of retained dimensions was further evaluated using Cattell’s scree test, whereby the scree plot was visually inspected to identify the point of inflection beyond which additional components contributed minimally to explained variance. Only dimensions preceding this inflection point were retained for subsequent analyses.

Following initial dimension selection, the quality of representation (cos²) and the contribution of each behavioural variable to the retained dimensions were assessed. Variables showing low cos² values, weak contributions, or limited biological relevance were excluded in an iterative manner, and PCA models were re-run until a stable and interpretable dimensional structure was obtained. This stepwise process ensured that the final PCA included only variables that meaningfully contributed to the behavioural structure of each test.

Following selection of the number of retained dimensions, a Varimax Rotation (VR) was applied to enhance interpretability. This rotation redistributes factor loadings to minimize cross-loadings and ensure that variables load more strongly onto a single dimension. When visualized, closely related variables cluster more distinctly, facilitating interpretation. The names and extremes of the dimensions were subsequently determined based on the variables loading most strongly onto each rotated dimension.

### 3.4. Statistical Tests Used for Intra- and Inter-testing

Normality of the data for all tests was assessed through visual inspection of QQ-plots, which indicated deviations from normality. As a result, Spearman’s rank correlation was selected for all statistical comparisons due to its suitability for analyzing personality data. As a non-parametric test, it does not assume normality or linearity, making it ideal for ordinal data (e.g., reaction scores) and transformed measures (e.g., PCA scores) that may not meet parametric assumptions. In the context of this study, the key consideration is whether a cow with a higher rank in one test consistently ranks similarly across tests, indicating intra- or inter-test consistency. Spearman’s method ranks the data rather than using raw values, enabling comparisons across different scales by focusing on the relative positioning of individuals within a group, rather than exact numerical differences. This is especially important given the varying scales across tests. Additionally, Spearman’s correlation identifies monotonic relationships, capturing trends where variables move in the same or opposite direction, revealing patterns of relatedness and consistency in the traits compared.

Statistical significance for all tests was defined at a p-value ≤ 0.05, with trends noted for p-values between 0.05 and 0.10. When correlations were significant, the strength of the relationships (ρ) was interpreted as follows: 0.00–0.19 = very weak or no relationship; 0.20–0.39 = weak; 0.40–0.59 = moderate; 0.60–0.79 = strong; and 0.80–1.00 = very strong relationship (Evans, 1996; Papageorgiou, 2022).

### 3.5. Intra-Test Consistency for Scored Tests and Software-Annotated Tests

Consistency over time is a key aspect of personality; therefore, Spearman’s rank correlation was employed to evaluate correlations across the three testing periods. For the Human Approach Test (HAT), additional comparisons were made between different stages within each period to account for the different locations of the human and the distinct actions performed in relation to the cows.

For the three behavioural tests with PCA graphs, individual coordinates from each test were extracted and compiled into data frames for intra-test comparisons. Individuals with data from only one period were excluded, as intra-test consistency analysis requires at least two periods for pairwise comparisons. Therefore, cows that completed two or three testing periods were retained in the analysis, allowing pairwise comparisons between available periods.

### 3.6. Inter-test Consistency Between Behavioural Tests

#### 3.6.1. Scored Tests (HAT and SUD)

Although conducted in different contexts, both the HAT and the SUD may evaluate related traits (e.g., boldness and fearfulness in response to stimuli). Examining correlations between variables measured in these tests can reveal whether cows that display confidence or high reactivity in one test exhibit similar behavioural tendencies in the other, indicating the presence of a shared underlying personality trait. Significant correlations would further suggest consistency of behaviours across various situations, reinforcing the reliability of these tests for assessing personality traits. Therefore, inter-test consistency was evaluated by comparing HAT and SUD results within corresponding periods (e.g., Period 1 HAT and Period 1 SUD).

#### 3.6.2. Software-Analyzed Tests (HAT, OFT, NST)

Evaluating inter-test consistency between PCA-derived behavioural tests requires consideration of several key factors. First, if intra-test consistency shows no significant correlations across periods, this suggests that the test may not reliably capture stable behavioural traits under the conditions tested, or that it may not be optimally suited to address the specific research question of identifying stable personality traits in dairy cows. Since personality is defined by behavioural responses that remain relatively stable over time and across contexts (Réale et al., 2010), a lack of consistency over time undermines the test’s validity.

Second, if none of the three tests exhibit consistency over time, inter-test correlations are unlikely to be meaningful, as unstable measures cannot reliably capture underlying personality traits.

However, since personality consists of both consistency over time and across situations, it may be valuable to compare the first period of each test to assess whether consistency exists at the contextual level, particularly if no correlation is observed between periods within each test (suggesting a lack of consistency over time). The first period of each test was chosen because it generally reflects the cow’s natural, unaltered state, less likely influenced by prior exposure or adaptation to the testing environment. This period provides a clear baseline for understanding a cow’s behaviour during its initial encounter with the test conditions. By focusing on this first period, potential biases from repeated testing or learning effects are minimized, ensuring that the observed behaviours more accurately reflect the cow’s innate responses. This approach enables assessment of personality consistency within each test’s context, even when traits vary over time, and reveals whether a cow’s behaviour in one test aligns with its behaviour in another.

To avoid making assumptions about the personality dimensions associated with each test, all possible inter-test comparisons were conducted, with the results revealing whether the two tests measure similar personality traits within the same dimension. For example, if two dimensions from different tests are positively correlated, it would suggest that cows with high scores in one dimension of one test (e.g., nervous response style in the Novel Object Test) are likely to also exhibit high scores in the corresponding dimension of another test (e.g., active transitioning in the Novel Surface Test). However, meaningful interpretation of inter-test correlations requires that the dimensions being compared are expressed on comparable behavioural scales. When dimensions differ fundamentally in their underlying behavioural constructs, observed associations may reflect differences in measurement structure or test design rather than shared personality traits, indicating that the tests or dimensions may not be optimally suited to address the question of whether different behavioural tests capture comparable personality traits. For instance, a scale from neutral (low reactivity) to extreme (high reactivity) cannot be compared to another scale that ranges from one extreme (cautiousness) to another (curiosity), as this could incorrectly suggest that a cow with lower reactivity (e.g., resting or feeding behaviours) is associated with cautiousness. Using comparable scales ensures that observed differences reflect true behavioural variation, rather than discrepancies in measurement or terminology. In some cases, data transformation may be required when two dimensions share a similar relative scale but are inversely related. In these cases, all values of one dimension will be multiplied by −1 to align the high scores across both datasets, ensuring they reflect the same behavioural tendencies. Standardizing these dimensions guarantees that any negative correlation represents a genuine behavioural difference rather than an artifact of differing scales.

## 4. RESULTS

### 4.1. Intra and Inter-Test Consistency of Scored Tests

This section presents the results of both intra- and inter-test reliability analyses based on scores from the Human Approach Test (HAT) and the Sudden Test (SUD). The intra-test analysis evaluated the consistency and stability of scores in relation to behavioural expressions over time within a single test, while the inter-test analysis examined overlapping traits, with significant positive correlations indicating that cows displaying similar behaviours in one test tended to respond comparably in the other.

#### 4.1.1. Intra-Test Consistency for the Human Approach Test (HAT)

Significant positive correlations in behavioural response scores were found between Periods 2 and 3 (ρ = 0.34, p-value = 0.02, df = 42). No significant correlations were observed between Periods 1 and 2 (ρ = 0, p-value = 1, df = 42) or Periods 1 and 3 (ρ = 0.18, p-value = 0.25, df = 42). When comparing the different stages of the test, no significant correlations were found in Stage 1 (Period 1 and 2: ρ = −0.05, p-value = 0.76; Period 2 and 3: ρ = −0.02, p-value = 0.88; Period 1 and 3: ρ = −0.05, p-value = 0.76). In Stage 2, a negative correlation was significant between Period 1 and 2 (ρ = - 0.31, p-value = 0.04), while a positive tendency was observed between Period 2 and 3 (ρ = 0.27, p-value = 0.08); no significant correlation was found in Period 1 and 3 (ρ = 0.09, p-value = 0.58). Stage 3 showed no significant correlations (Period 1 and 2: ρ = 0.04, p-value = 0.78; Period 2 and 3: ρ = 0.21, p-value = 0.17; Period 1 and 3: ρ = 0.17, p-value = 0.27). In Stage 4, two pairwise comparisons exhibited positive correlations (Period 2 and 3: ρ = 0.43, p-value = 0.004; Period 1 and 3: ρ = 0.49, p-value < 0.001), while one comparison showed no significance (Period 1 and 2: ρ = 0.24, p-value = 0.11). For Stage 5, one pairwise comparison was significant (Period 2 and 3: ρ = 0.57, p-value < 0.001), with a positive trend observed between Period 1 and 2 (ρ = 0.25, p-value = 0.09), but no correlation was found between Period 1 and 3 (ρ = 0.16, p-value = 0.29). The degrees of freedom for all stages was 42.

Correlations per stage are reported here but will not be used in the HAT analysis for the remainder of the paper for two main reasons: (1) no consistent correlations were found between stages, and (2) the analysis focused on overall behavioural patterns across the entire testing period, rather than responses within individual stages.

#### 4.1.2. Intra-test consistency for the Sudden Test (SUD)

Significant positive correlations were found between Periods 1 and 2 (ρ = 0.47, p-value < 0.001) and between Periods 1 and 3 (ρ = 0.37, p-value = 0.009). A trend toward significance was observed in the comparison between Periods 2 and 3 (ρ = 0.27, p-value = 0.06). The degrees of freedom for the three comparisons was 46.

#### 4.1.3. Inter-test consistency between the Human Approach Test (HAT) and the Sudden test (SUD)

When comparing between the three periods for each test (e.g., Period 1 HAT vs. Period 1 SUD), significant positive correlations were found across all three pairwise period comparisons (df = 42): Period 1 (ρ = 0.60, p-value < 0.001), Period 2 (ρ = 0.41, p-value = 0.01), and Period 3 (ρ = 0.43, p-value = 0.008).

### 4.2. Principal Component Analysis (PCA) of Software-Annotated Behavioural Tests

In this section, the PCA dimensions identified through the data manipulation strategies outlined in the *3. Statistical Analysis* section will serve as the foundation for identifying potential personality traits associated with each test. These dimensions will also serve as a model for assessing both intra- and inter-test consistency, as the PCA results will reveal the cows’ coordinates (indicating their position at each period) in relation to the two dimensions.

#### 4.2.1. Novel Object Test (NOT) PCA

The PCA identified two behavioural dimensions in cows’ responses to novel objects, both with eigenvalues greater than one. Together, these dimensions accounted for 54.33% of the variance (Figure 7).

**Figure 7.**
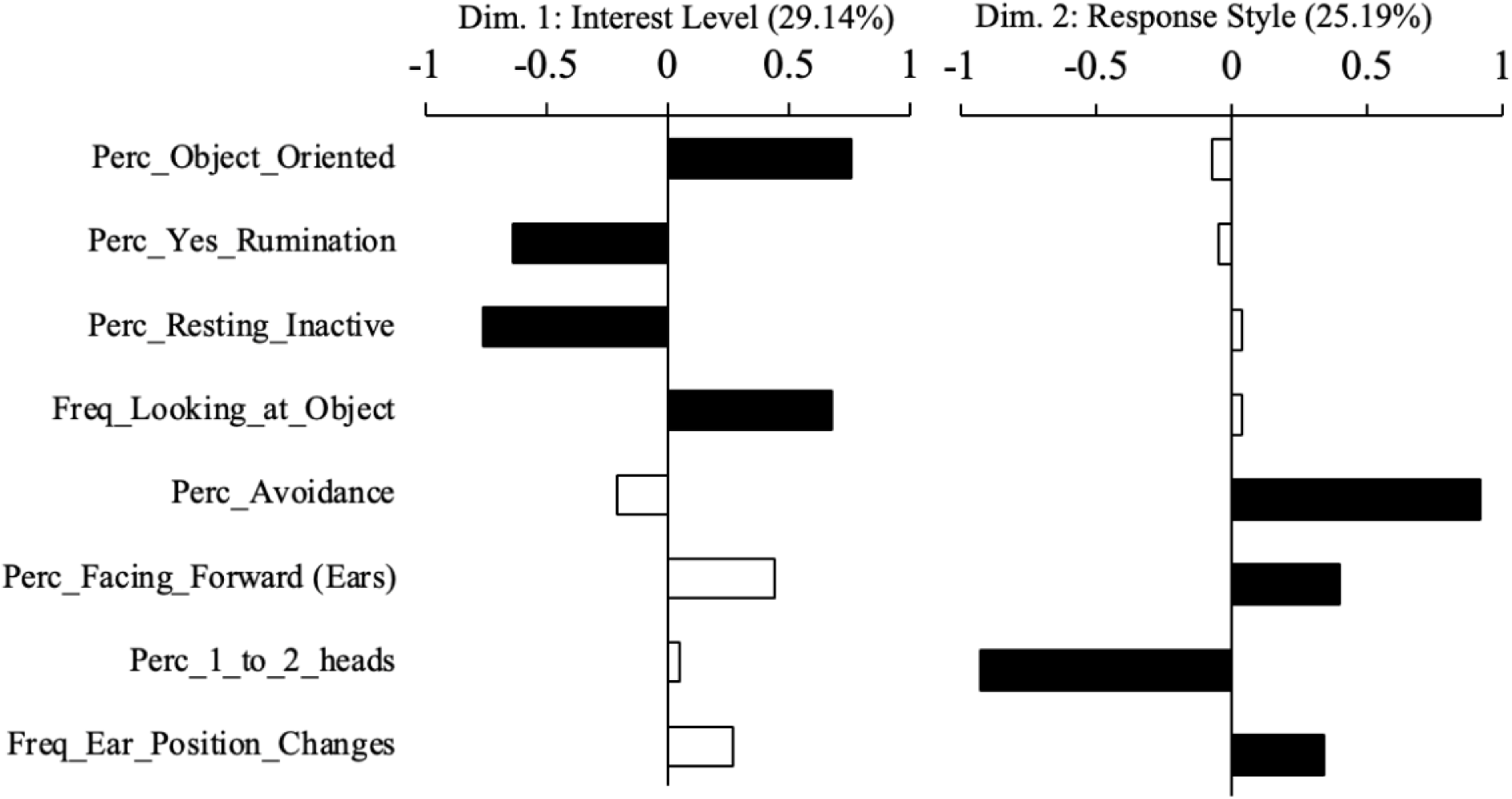
Coordinates of variables from the PCA, mapped onto Dimension 1 (Interest Level, ranging from passivity to curiosity) and Dimension 2 (Response Style, ranging from neutral to nervousness). Variables with higher contributions to each dimension are represented by black bars, indicating their stronger influence within the respective dimension.

The first dimension, labeled “Interest Level,” ranged from passivity to curiosity. It encompassed behaviours such as resting, inactivity, and rumination, which opposed behaviours like frequently looking at the novel object and engaging with it through licking, sniffing, biting or pushing.

The second dimension, labeled “Response Style,” ranged from neutral to nervousness. This dimension included the percentage of time spent within 1–2 head lengths away from the object, contrasted by behaviours such as frequency of ear position changes, the proportion of time with ears facing forward, and time spent avoiding the object. Avoidance behaviours were characterized by maintaining a distance greater than two head lengths, stretching the neck chain to its limit, or standing up/moving away in response to the object.

#### 4.2.2. Open Field Test (OFT) PCA

The PCA revealed two dimensions characterizing cows’ behavioural responses to isolation in a novel arena, both with eigenvalues greater than one, collectively explaining 49.68% of the variance (Figure 8).

**Figure 8.**
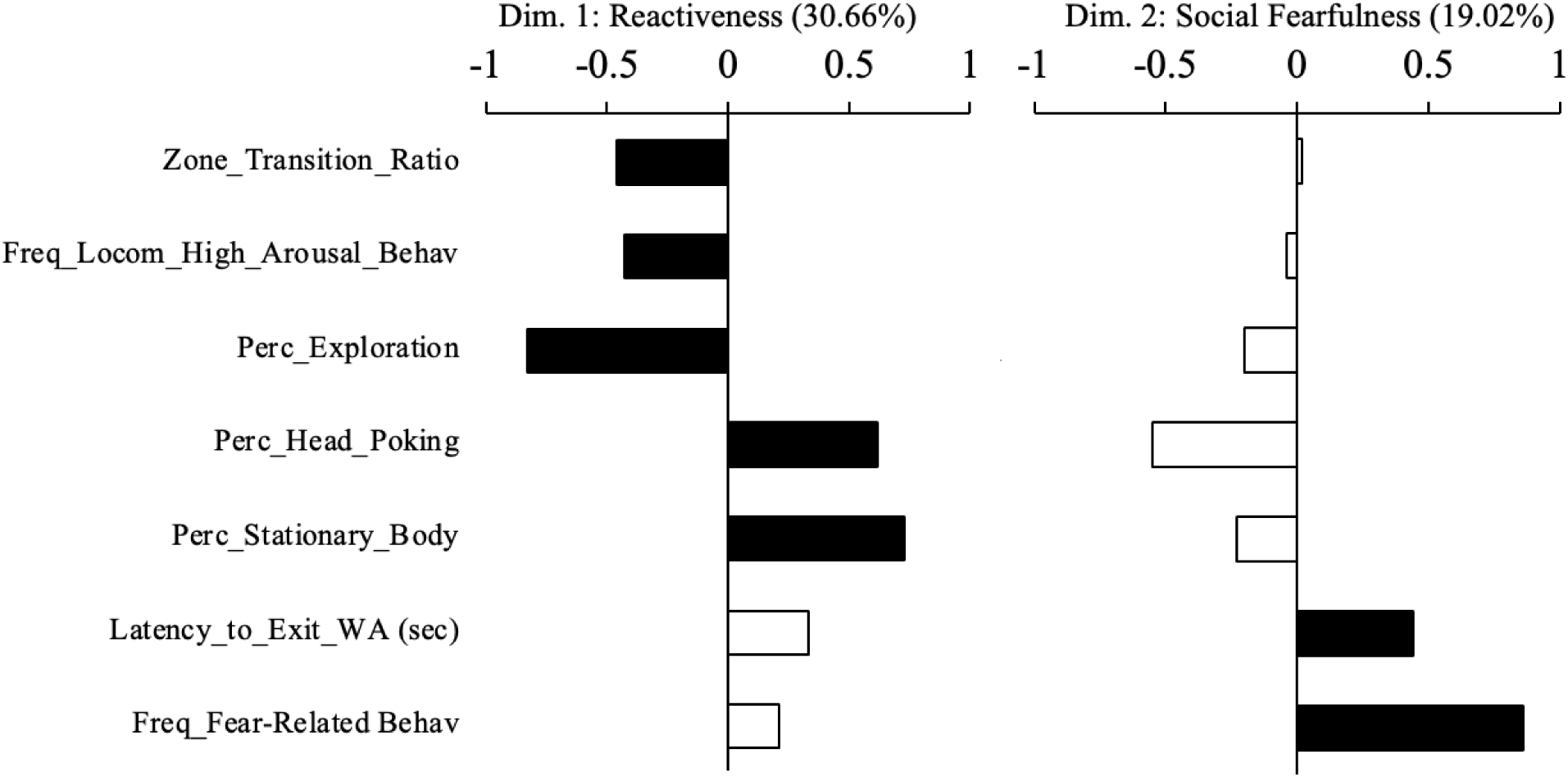
PCA variable coordinates mapped onto Dimension 1 (Reactiveness, ranging from high to low reactivity) and Dimension 2 (Social Fearfulness, ranging from comfort to fear in isolation). Variables with higher contributions to each dimension are highlighted with black bars, emphasizing their stronger influence within the respective dimension.

The first dimension, labeled “Reactiveness”, ranged from high to low reactivity. It encompassed behaviours such as zone transitions, exploration, and high-arousal locomotor actions (i.e., charging, trotting, and pacing), with head poking out of the arena and a stationary body positioned at the opposite extreme.

The second dimension, termed “Social Fearfulness”, spanned from comfort to fear in isolation. Since no variables were present on the opposite side of the “fear of isolation” extreme, which included indicators like jumping, vigilance, and vocalization, we infer that this side represents the inverse of fear, likely reflecting calm or neutral behaviours.

#### 4.2.3. Novel Surface Test (NST) PCA

The PCA identified two behavioural dimensions in cows’ responses to a novel surface in a corridor, both with eigenvalues greater than one, collectively explaining 71.75% of the variance (Figure 9).

**Figure 9.**
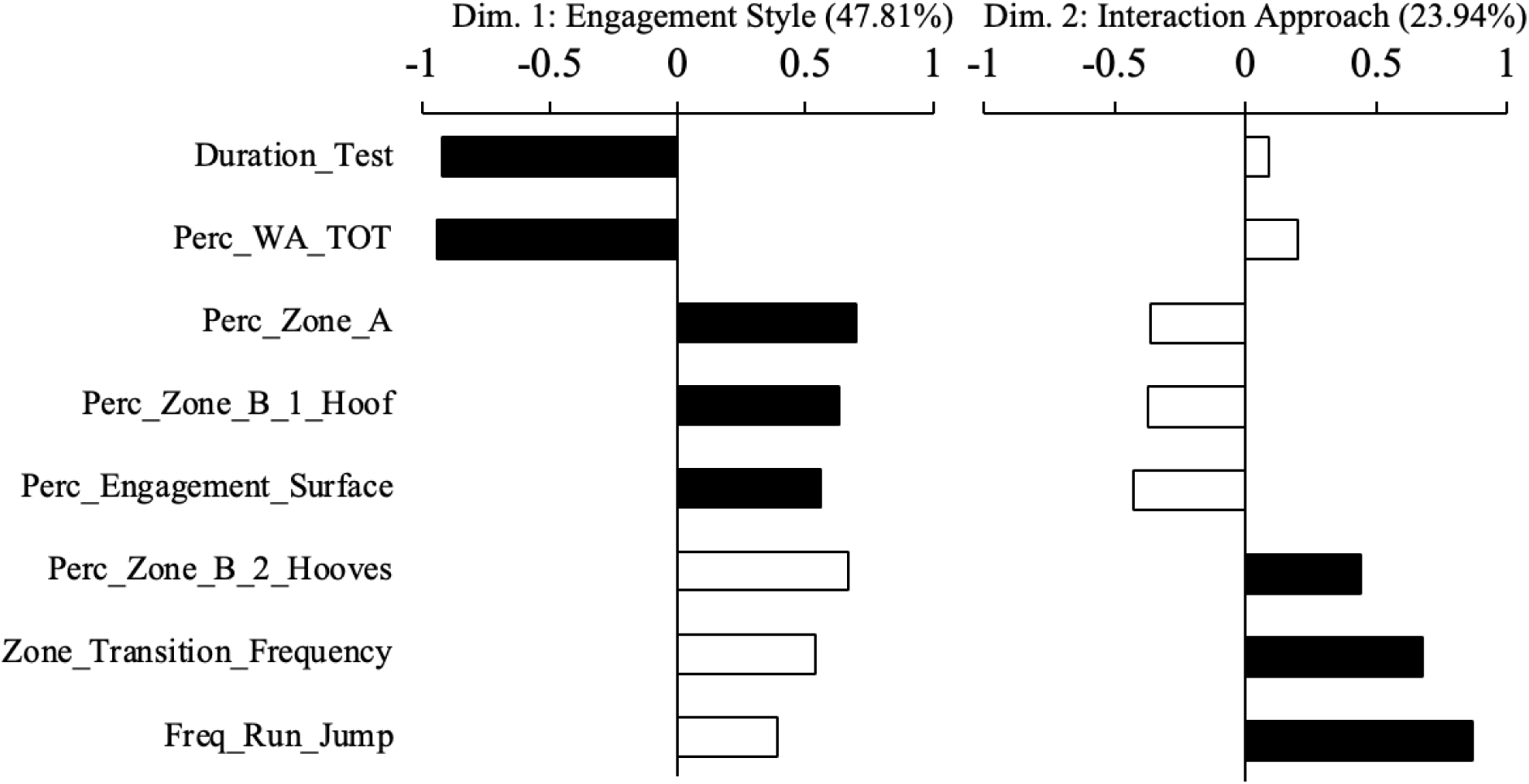
PCA variable coordinates plotted along Dimension 1 (Engagement Style, spanning from cautiousness to curiosity) and Dimension 2 (Interaction Approach, ranging from inactivity to active transitioning). Black bars highlight variables with higher contributions to each dimension, underscoring their greater influence within the respective dimension.

The first dimension, labeled “Engagement Style,” ranged from cautiousness to curiousness. This dimension included behaviours such as the percentage of time spent in the waiting area and the duration taken to complete the test, contrasted with the percentage of time spent with one hoof on the surface, time engaging with the surface (including exploration and foot tapping), and time spent in Zone A (the area closest to the surface).

The second dimension, termed “Interaction Approach,” ranged from inactive to active transitioning. No variables were present on the opposite side of the “Active Transition” extreme, which included behaviours such as the frequency of running, jumping, placing two hooves on the surface, and zone transitions. As a result, we inferred that this side represents the inverse of activeness, likely indicating stationary or inactive behaviours, and therefore labeled the opposite extreme as “Inactiveness.”

### 4.3. Intra-Test Consistency of Tests Using Principal Component Analysis (PCA)

This section presents the results of the intra-test consistency analysis based on the individuals’ PCA coordinates. This analysis evaluates the stability of cows’ positions along the dimensions over time, with significant correlations between periods indicating behavioural consistency. Such consistency would further support the validity of this test in measuring personality traits.

#### 4.3.1. Novel Object Test (NOT)

No significant correlations were observed for Dimension 1 (Interest Level) coordinates of the NOT across testing periods: Period 1 vs. 2 (ρ = 0.11, p-value = 0.55, df =33), Period 1 vs. 3 (ρ = 0.18, p-value = 0.30, df = 33), and Period 2 vs. 3 (ρ = 0.13, p-value = 0.40, df = 33). Likewise, Dimension 2 (Response Style) coordinates showed no significant relationships: Period 1 vs. 2 (ρ = −0.09, p-value = 0.60, df = 33), Period 1 vs. 3 (ρ = 0.14, p-value = 0.43, df = 33), and Period 2 vs. 3 (ρ = 0.07, p-value = 0.62, df = 45).

#### 4.3.2. Open Field Test (OFT)

For Dimension 1 (Reactiveness) coordinates of the OFT, no significant correlations were found between periods: Period 1 vs. 2 (ρ = −0.03, p-value = 0.91, df = 18), Period 1 vs. 3 (ρ = 0.10, p-value = 0.67, df = 20), and Period 2 vs. 3 (ρ = 0.29, p-value = 0.17, df = 22).

For Dimension 2 (Social Fearfulness) coordinates, a moderate positive correlation (ρ = 0.61, p-value = 0.01, df = 18) was found between Period 1 and 2. However, no significant correlations were detected for the other comparisons: Period 1 vs. 3 (ρ = −0.21, p-value = 0.35, df = 20) and Period 2 vs. 3 (ρ = 0.06, p-value = 0.76, df = 22).

#### 4.3.3. Novel Surface Test (NST)

For Dimension 1 (Engagement Style) coordinates of the NST, no significant correlation was found between Period 1 and 2 (ρ = 0.34, p-value = 0.29, df = 10). However, a moderate positive correlation with a tendency towards significance was observed between Period 2 and 3 (ρ = 0.45, p-value = 0.09, df = 13). A significant moderate positive correlation was detected between Period 1 and 3 (ρ = 0.55, p-value = 0.04, df = 13).

For Dimension 2 (Interaction Approach) coordinates, no significant correlations were found in any of the pairwise period comparisons: Period 1 vs. 2 (ρ = 0.28, p-value = 0.38, df = 10), Period 1 vs. 3 (ρ = 0.41, p-value = 0.13, df = 13), and Period 2 vs. 3 (ρ = 0.16, p-value = 0.56, df = 13).

### 4.4. Inter-Test Consistency of Tests Using Principal Component Analysis (PCA)

This section presents the results of the inter-test consistency analysis based on individuals’ coordinates across dimensions for different tests. This analysis aimed to identify shared traits between tests, where significant positive correlations suggest that cows displaying similar behaviours in one test are likely to exhibit comparable responses in the other.

No significant correlation was observed between the following pair-wise dimension comparisons identified based on similar scaling:

| Pairwise comparison between dimensions of PCAs | | Correlation coefficient ($\rho$) | P-value | Degrees of freedom (df) |
| --- | --- | --- | --- | --- |
| First Dimension | Second Dimension |  |  |  |
| | Dimension 1 NOT (**Interest level**) | $\rho = -0.03$ | p = 0.91 | 13 |
| Dimension 2 NST<br>(Interaction approach) | Dimension 1 OFT (**Reactivity**) | $\rho = 0.05$ | $p = 0.86$ | 11 |
| | Dimension 2 NOT (**Response Style**) | $\rho < 0.001$ | $p = 1$ | 13 |
| Dimension 1 OFT<br>(**Reactivity**) | Dimension 1 NOT (**Interest level**) | $\rho = 0.04$ | $p = 0.87$ | 9 |
| | Dimension 2 NOT (**Response Style**) | $\rho = -0.05$ | $p = 0.84$ | 18 |
| Dimension 1 NST<br>(**Engagement style**) | Dimension 2 OFT (**Social Fearfulness**) | $\rho = 0.29$ | $p = 0.33$ | 11 |

## 5. DISCUSSION

### 5.1. Intra-Test Consistency for Behavioural Tests

#### 5.1.1. Human Approach Test (HAT)

The correlation of individual responsiveness was assessed across three time periods with 1-month intervals. Only one pairwise comparison (Period 2 vs. Period 3) showed a weak but significant correlation. This suggests that cows may have responded more consistently once the novelty of the human presence and the testing procedures in general had diminished. While initial responses (in Period 1) may have reflected raw, unconditioned responses to the unfamiliar situation, cows in later periods may have become more accustomed to the testing context. However, this habituation effect may not have been solely due to repeated exposure to the same test but could also reflect the cumulative experience of being confronted with multiple novel situations within a short time frame, including a new human, a novel object, a novel surface, and an unfamiliar environment.

The amount of human handling and prior experience may have also influenced the cows’ behavioural responses; however, it is likely that these factors did not affect all individuals in the same way. Previous studies suggest that approach behaviours towards humans can be driven by various emotions or motivations, such as curiosity towards novel stimuli (e.g., an unfamiliar human; Welp et al., 2004) or positive expectations associated with the approach and human contact (e.g., brushing or stroking, Jago et al., 1999), while fear may prompt the avoidance of the stimulus (Breuer et al., 2000). In this context, variability in prior experiences or sensitivity to novelty may have influenced individual responses over time, leading to changes in rank order consistency between Period 1 and later periods. Moreover, because many of the cows were routinely exposed to humans for research purposes and were tested in their home environment, the perceived threat or novelty of the human may have decreased more rapidly, resulting in less pronounced behavioural variation. Supporting this, Doyle et al. (2022) found that dairy calves tested in their home pens during a human approach test scored higher, indicating greater proximity to the human, than those tested in a novel arena. This highlights how environmental familiarity (and potentially the presence of conspecifics) can contribute to a group-level calming effect, further reducing individual variation in responses.

Unlike previous studies that used very short intervals between tests (e.g., 2-day intervals in Lensink et al., 2003; 3–4 days in Gibbons et al., 2009), the present study adopted a one-month interval between tests. This approach aimed to balance two key considerations: preserving the unfamiliarity of the human stimulus while maintaining a relatively stable testing context. Shorter intervals increase the risk of habituation, whereas longer intervals, such as those used in studies comparing calves to adult cows or using 8-month gaps (Hasenpusch et al., 2024), may introduce greater variability in factors such as age, parity, or days in milk. The one-month interval chosen here was intended to provide a middle ground for test validation within a comparable context.

Nonetheless, even within this timeframe, certain changes in the broader testing environment could not be entirely controlled. Routine management practices, such as stall repositioning, regrouping, or shifts in outing schedules, alongside environmental factors like increasing temperatures in later months that can induce greater lethargy (Hansen & Aréchiga, 1997), may have influenced cows’ behavioural responses. However, it is not necessarily the environmental change itself that compromises consistency, but rather the potential for individual differences in how cows cope with or adapt to these changes. In our case, consistency is evaluated based on the stability of individual ranks relative to others, rather than on absolute behavioural scores, a methodological approach similarly employed in previous work (Zambelis & Vasseur, 2022). Therefore, if cows responded uniformly to contextual changes (e.g., all becoming more lethargic in hotter conditions), rank order consistency would likely be maintained. In contrast, variable individual responses to such changes may have contributed to weaker correlations observed between certain periods in this study.

This raises a broader concern regarding the limitations of tests like the HAT in consistently measuring personality. The loss of novelty over time, due to habituation or environmental factors, may undermine the test’s ability to accurately assess stable personality traits. As personality is often defined by relatively stable behaviours across contexts and time (Réale et al., 2010), repeated exposure to the same testing environment could introduce significant variability in responses, making it difficult to measure consistent individual traits. If the test itself becomes too familiar or predictable, the behaviours observed may no longer reflect inherent personality traits but instead reflect the cows’ changing emotional responses to the testing situation, such as fatigue, stress, or prior experiences, rather than inherent personality traits.

Therefore, while the HAT provides valuable insight into how cows respond to human interaction, its reliability for assessing consistent personality traits may be limited by the loss of novelty over repeated exposures. In contrast to our study, Hasenpusch et al. (2024) evaluated the consistency of personality traits using a Forced Human Approach Test (FHAT—equivalent to the HAT used here) and found minimal differences between test and retest scores, suggesting higher consistency. However, their study used an 8-month interval between tests, during which cows may have had limited exposure to novel situations, potentially preserving their sensitivity to novelty at retest. In contrast, cows in our study were tested monthly over three consecutive months, a shorter interval that may have increased their general familiarity with being confronted by novel situations or test procedures, particularly given the research context where cows were often handled or exposed to experimental setups. Moreover, previous findings suggest that dairy cows can retain memory of individuals or experiences for several weeks (Forkman et al., 2007), which may have further influenced their responses. Additionally, individual differences in habituation rates or coping styles may have further contributed to the variability in responses observed over time.

Additionally, it is worth noting that a set of unidimensional “responsiveness” score was used to quantify the magnitude of behavioural reactions, whether positive (e.g., approach behaviours) or negative (e.g., retreating behaviours), to the human stimulus. This approach was chosen to capture individual variation in the intensity of behavioural engagement, irrespective of its direction. The rationale for this decision is grounded in the assumption that animals displaying stronger reactions in either direction may possess a more reactive or emotionally responsive temperament, while those with weaker or absent responses may be less emotionally engaged or more passive. According to Rothbart and Derryberry (1981), an individual’s temperament, which is considered to be a subset of the broader domain of individual differences in personality, reflects biologically based individual differences in emotional reactivity and self-regulation, which can be measured through the intensity and duration of responses. Individuals displaying stronger emotional or behavioural reactions may be considered more reactive or emotionally responsive, while those showing weak or absent responses may possess lower emotional engagement or passivity. While this approach may conflate motivational valence (i.e., whether the animal is experiencing the situation as positive or negative), it offers a pragmatic means of identifying individuals who are more behaviourally expressive overall. Such expressiveness may reflect their general emotionality or sensitivity to stimulation, traits that can emerge in both negative and positive situations.

However, it is important to recognize that this simplification comes with limitations. Pooling approach and avoidance behaviours may obscure meaningful distinctions between different personality dimensions, such as openness to experience (expressed through boldness/shyness towards the human) versus neuroticism (fearfulness). Future analyses could benefit from separating directional valence and intensity of responses, or from applying a multidimensional scoring approach that could distinguish between the individual’s motivation and their arousal level. Despite this limitation, the unidimensional approach was selected here for its practical utility in summarizing behavioural variability, and for facilitating comparisons across multiple tests and timepoints.

To validate the use of the HAT or similar tests, future studies should aim not only to refine the methodology but also to address the impact of novelty loss when retesting occurs over shorter intervals. While short intervals between tests may be better suited for validating the test itself by controlling for context and ensuring that repeated measures yield consistent results under similar conditions, longer intervals may be more appropriate for validating personality traits over time. Testing over longer intervals introduces natural changes in context (e.g., age, parity, season), allowing researchers to assess whether individual differences remain stable despite these changes, which is a core criterion for personality assessment.

#### 5.1.2. Sudden Test (SUD)

The correlations observed for the SUD suggest that this test may be a useful tool for measuring stable personality traits in dairy cows. Consistency was observed across all pairwise comparisons of testing periods, with the strongest relationship found between the first two periods, and weaker but still notable relationships in later comparisons. While the correlation between Periods 2 and 3 did not reach the p < 0.05 threshold, the overall pattern supports the potential of the SUD as a relatively reliable measure of individual behavioural tendencies over time. In contrast to other behavioural tests evaluated in this study, the SUD captures immediate, unconditioned reactions of the cows, rather than behaviours shaped by prior learning or adaptation to repeated stimuli. Previous studies have suggested that bolder animals may learn novel situations more quickly and adjust their behaviour as they become familiar with the task (Yuen et al., 2017). For instance, in an Open Field Test, once a cow enters the testing arena and waits in the start box, she may rely on indirect environmental cues to assess potential risks and adjust her coping strategies accordingly (Yuen et al., 2017). As the cow progresses through later periods (2 and 3), prior exposure to the arena allows her to modify her behaviour to better handle the situation (Frost et al., 2006). However, in the SUD, no preparatory cues are provided, so the cows’ immediate reactions to the suddenness of the stimulus are more likely to reflect their true personality traits and remain consistent over time. However, certain methodological limitations should be considered. Occasional malfunctions of the pulley system may have resulted in objects dropping at a speed insufficient to induce a fear response. Additionally, some cows may not have noticed the sudden appearance of the object if it was outside their peripheral vision or if they were distracted at the time (e.g., by feed). These factors could have contributed to variability in individual reactions, potentially affecting the reliability of the test.

#### 5.1.3. Novel Object Test (NOT)

No significant correlation was found when comparing each period for individual cows across the two PCA dimensions (Interest Level and Response Style), indicating that behaviours expressed in one period were not consistently carried over to subsequent periods. Several factors may contribute to this lack of repeatability.

First, unlike the SUD, which captures an individual’s immediate reaction to a sudden stimulus, the NOT requires cows to notice, process, and decide how to interact with the novel object. However, within the five-minute testing window, external distractions (e.g., hunger, fatigue, presence of conspecifics, or environmental stimuli such as heat that place self-regulation above other interests) may have diverted their attention away from the object, leading to inconsistent behavioural responses.

Second, to control variability in responses to the novel object, we ensured that its texture remained constant while only the shape changed across periods. This approach aimed to minimize potential biases arising from individual preferences for certain textures, which could otherwise lead to differences in exploratory behaviour. However, while this reduced one source of variation, the factors that make an object engaging or intimidating remain largely unexplored in current literature. The response to a novel object is likely influenced by multiple properties, including size, odor, shape, color, and potential resemblance to familiar objects, all of which can impact whether the object is perceived as neutral, interesting, or threatening (Takola et al., 2021).

As a result, behavioural responses may vary not only due to the specific features of the object itself but also based on individual differences in how cows perceive and interpret the object in that particular period. If an object is seen as intimidating, the cow may exhibit freezing, avoidance, or fear-related behaviours, whereas a neutral or engaging object may promote exploration and interaction (Hughes, 1997). For example, if the provided stimuli do not result in or lead to a threat (e.g., restraint after introduction of novelty [Baugh et al., 2017]) or a reward (e.g., objects that provide grooming opportunities [Dickson et al., 2024]), the cow’s reactions may diminish over time, as there is neither a motivation to approach nor a need to avoid the object, leading to inconsistent responses. While we controlled for texture to reduce variation due to tactile preference, the objects still differed in shape across periods to maintain novelty. It is possible that some shapes were perceived as more neutral, while others resembled threatening or overly familiar stimuli, leading to varied reactions depending on the cow’s interpretation of each object. This variation in perception, despite consistent material, could have contributed to the lack of behavioural consistency observed in the NOT. Therefore, even subtle changes in object shape may influence how novelty is perceived, and future studies might benefit from standardizing both material and shape, or systematically testing how specific object features (e.g., symmetry or size) impact cow behaviour.

Additionally, as suggested by previous studies, whether an individual is tested with or without social isolation can be a critical factor in the design of the NOT. The reactions of social animals to novelty can be influenced by the presence of conspecifics, as social interactions may modify behavioural expressions (Hirata et al., 2016). For instance, kinship or familiarity between individuals can reduce the likelihood of aggression, while the lack of relatedness may induce stress (Bartolomucci et al., 2002), potentially leading to less exploratory behaviours and more avoidance (Matisz et al., 2021). While some cows were moved to new stalls for regrouping purposes between testing periods, these changes were made well in advance of the tests to minimize disruption, stabilize social dynamics, and reduce any potential impact on behavioural expression across periods.

Importantly, we chose to conduct testing within the home pen rather than in social isolation, as removing cows from their social group could have introduced an even greater stressor, potentially overshadowing their response to the novel object itself. Although testing within the group comes with some variability, this approach likely reflects a more ecologically valid representation of their behavioural tendencies. In future work, maintaining more consistent group compositions across testing periods or assessing individual responses in a semi-isolated but familiar setting may help to further refine behavioural consistency.

Despite these challenges, our results raise important questions about the reliability of the NOT for assessing stable personality traits in adult dairy cows. While this test has increased in popularity in animal personality research (Takola et al., 2021), perhaps for its simplicity and non-invasive nature, the lack of consistent behavioural patterns across testing periods in our study and others (e.g., Kilgour et al., 2006; Lansade et al., 2007) suggests that it may not be a robust standalone measure of personality over time. However, contrasting evidence highlights the complexity of this issue. For instance, Hasenpusch et al. (2024) reported consistent trait expression between test and retest using the NOT and Forced Human Approach Test (FHAT) with an 8-month interval, suggesting that the detection of stable traits may depend on various factors, such as test interval, context, or prior experience. This is particularly relevant for dairy cows, whose attentional focus, motivational state, and interpretation of novelty may shift subtly across time and situations, possibly influencing test outcomes.

Our study contributes uniquely to this discussion by systematically testing the NOT across multiple periods and controlling for object material, a variable not consistently addressed in previous work (Knauer et al., 2021; Amy et al., 2017; Hirata et al., 2018). While we observed variability in behaviour, the fact that this inconsistency persisted despite efforts to standardize key elements (e.g., object texture, familiar environment) suggests that the test itself may be overly sensitive to transient factors. This could reflect a limitation in its design when used as a longitudinal tool to assess personality.

Therefore, although the NOT may still hold value as a single-timepoint measure to quantify shyness–boldness, exploration–avoidance or neophilia/neophobia in an individual (Takola et al., 2021), its usefulness in identifying consistent, long-term personality traits in dairy cattle appears limited. Future research might explore adaptations to the NOT, such as extending the exposure time or introducing motivational outcomes (e.g., reward or mild challenge), to better capture stable behavioural traits.

#### 5.1.4. Open Field Test

No significant correlation was found in the pairwise comparisons of PCA Dimension 1 (Reactiveness) between all 3 periods, but one significant strong positive correlation was observed in Dimension 2 (Social Fearfulness) between Periods 1 and 2. The remaining period comparisons were not significant for Dimension 2. This suggests that in terms of social fearfulness displayed by the cows, their comfort or fear within the isolated arena remain relatively constant over the 2 time periods, though it did not persist to the 3^rd^ period. Cows are naturally gregarious animals (Keeling & Gouyou, 2001), meaning they have an inherent social structure and prefer to be in the company of other cows. As such, their response to isolation may remain relatively consistent over time. This could explain why the correlation for social fearfulness between Periods 1 and 2 was moderately significant, indicating that the cows’ response to being isolated remained stable. Given that social fearfulness is likely tied to their need for social interaction and their natural discomfort with isolation, it is reasonable to expect that cows would consistently exhibit comparable levels of fear or discomfort when isolated from their group.

However, the lack of consistent results across all three periods raises uncertainty regarding the OFT’s reliability as a longitudinal measure of personality. While a single significant correlation suggests that some traits, like social fearfulness, may be expressed consistently in the short term, the absence of consistency across all time points limits the strength of this conclusion.

Additionally, like other tests involving exposure to novel situations, the OFT is susceptible to habituation effects. While personality is generally considered to be stable over time, the behavioural expressions of personality traits, particularly those associated with reactivity to novelty (such as shyness), can be influenced by repeated exposure to the same environment (Drent et al., 2003; Koolhaas & Van Reenen, 2016). As such, it is important to account for habituation when interpreting behavioural responses in repeated tests (Perals et al., 2017). In this context, the dimension of reactiveness may have been particularly sensitive to habituation. Some consistency in behaviour was observed across periods, especially in response to unpredictable stimuli, as seen in the SUD. However, because the same arena was used across the repeated OFT sessions, it is possible that the cows became familiar with the setting over time, which may have affected how strongly their underlying traits were expressed. The OFT was chosen over the Novel Environment Test for its ability to assess emotionality, fear, boldness, and exploration in individuals (Walsh & Cummins, 1976), offering a more comprehensive evaluation of animal personality compared to the Novel Environment Test, which introduces a new environment with each exposure but tends to focus on less personality-relevant factors, such as activity levels (Butler et al., 2011) and exploration-avoidance tendencies (Dingemanse et al., 2007). Although the one-month interval between testing periods was expected to help preserve the novelty of the arena, repeated exposure to the same environment may still have influenced the cows’ responses. For instance, an observed increase in escape attempts (results not presented) in Periods 2 and 3 suggests the cows may have learned the location of the exit and become comfortable enough to explore beyond or even leave the arena. This shift could reflect habituation to the context, rather than a change in personality itself.

Taken together, these findings suggest that while the OFT may be effective for capturing short-term responses to novelty or isolation, its ability to measure stable personality traits over time is limited in this study. This aligns with other work questioning the validity of the OFT as a personality test, although in this case the concern relates to a different aspect: specifically, its tendency to conflate multiple behavioural traits such as shyness, activity, and exploration within a single dimension (Perals et al., 2017). In the present study, it is possible that behaviours contributing to one dimension (e.g., Reactiveness) were not consistent because they reflect different underlying motivations or emotional states at different time points. For example, entering the novel arena may indicate boldness on one occasion but curiosity on another. These nuances suggest that repeated exposure to the same environment may not consistently activate the same trait. Furthermore, the lack of independent validation for which traits the OFT is measuring, as raised in previous research (Walsh & Cummins, 1976; Carter et al., 2012), reinforces the need to interpret trait labels and dimensions with caution.

#### 5.1.5. Novel Surface Test (NST)

No significant correlation was observed across periods for Dimension 2 (Interaction Approach), suggesting that cows did not maintain a stable tendency to be more active or inactive in their approach behaviours over time. In contrast, Dimension 1 (Engagement Style) showed more promising indications of stability. Although no consistency was evident between the first two periods (Periods 1 and 2), the pattern of correlations between later periods (Periods 2 and 3, Periods 1 and 3) suggests that cows’ tendencies to be more cautious or curious toward the novel surface may have remained relatively consistent over time.

A possible explanation for the lack of consistency observed for Dimension 2 may be attributed to varying levels of novelty across the different periods. Similar to the NOT, the NST also introduces individuals to novel stimuli. However, to our knowledge, no standardized method exists for balancing novelty, as animals’ responses to novelty are likely influenced by various properties, such as color, pattern, and texture (Takola et al., 2021). As a result, cows may have responded more strongly to certain surfaces, leading to more active behaviours such as running, jumping, and making more zone transitions. These behaviours could either suggest exploratory tendencies or heightened agitation or fear (de Passillé et al., 1995).

Similarly, for the first dimension of Engagement Style, the lack of behavioural consistency between Periods 1 and 2, but the presence of consistency in other pairwise comparisons, may suggest that cows exhibited a preference or curiosity for certain surfaces. For example, the carpet used in Period 2 with irregular patterning may have introduced a stronger visual novelty compared to the more structured patterns used in Periods 1 and 3 (grip tapes with equal spacing in Period 1 and checkered yoga mats with uniform areas in Period 3). Additionally, the softer surfaces, such as the carpet and yoga mats used in Period 2 and 3 respectively, may have resembled the rubber mats found in the cows’ stalls, leading to more comparable responses in those periods. In contrast, the grip-taped wood used in Period 1 was notably harder, introducing a distinctly different tactile experience.

Additionally, the design of the test involved placing two familiar companion cows on the far side of the novel surface. This setup meant that crossing the surface not only reflected a cow’s response to novelty, but also their motivation to rejoin social companions. As such, behaviours observed in this test may reflect an interaction between exploratory drive and social motivation, two traits that are relevant for both personality assessment and real-world group dynamics. However, this also complicates the interpretation of behaviours like latency to cross or zone transitions, since they may have been driven by differing combinations of social interest, fear, or boldness depending on the surface presented.

From an evaluative standpoint, these findings suggest that the NST, while promising for capturing short-term variation in exploratory behaviour, may not reliably measure stable personality traits unless the novelty component can be standardized. This aligns with challenges reported in similar novelty-based tests like the NOT, where the novelty type and properties can influence how cows perceive and respond to the test (Takola et al. 2021). To our knowledge, no published studies have yet validated the NST as a consistent personality test in dairy cattle across time. Our results add to this gap by showing that although certain dimensions (like Engagement Style) showed moderate temporal stability, others did not, likely due to the influence of stimulus properties.

In this sense, while the NST may not meet the criteria for a robust longitudinal personality test in its current form, it remains valuable for understanding context-dependent exploratory behaviour and social motivation. Further refinement, such as standardizing stimuli or separating the social and exploratory components, may improve its reliability as a personality assessment tool.

### 5.2. Inter-test Consistency for Behavioural Tests

#### 5.2.1. Human Approach Test (HAT) and the Sudden Test (SUD)

The inter-test results revealed significant moderate to strong correlations between the HAT and SUD when comparing the same testing period. This was somewhat unexpected, given that the HAT did not exhibit consistent behavioural responses across all periods. Typically, if a test lacks temporal consistency, one would expect weaker correlations with other tests, as measuring stable personality traits requires individuals to respond consistently across repeated exposures. Since HAT fluctuated over time while SUD remained relatively stable, their correlations might have been expected to be weaker, as individuals would not consistently score similarly across repeated testing.

To further explore these patterns, inter-test comparisons between the HAT and SUD were performed using data from all three testing periods. This approach differed from the inter-test comparisons performed for the NOT, OFT, and NST, which were limited to Period 1. The rationale for including all three periods for the HAT and SUD was based on the observation of some degree of temporal consistency across periods, particularly for the HAT, where the only significant positive correlation was found between Periods 2 and 3. This provided justification for performing inter-test consistency across all periods, as later periods still captured some stable individual differences. In contrast, for the NOT, OFT, and NST, no significant correlations were found between periods, suggesting a lack of temporal consistency. As such, comparisons for these tests were limited to Period 1, which was expected to reflect the cows’ most unconditioned and natural behavioural responses. If consistency between tests was to be found, it would most likely emerge during this initial exposure, prior to any potential habituation effects. Under this assumption, stronger correlations between the HAT and SUD might have been expected if both tests captured the same underlying traits. However, the moderate strength of these correlations may suggest that the two tests only partially overlap in what they assess, or that HAT responses are influenced by additional emotional or environmental factors, even during initial exposures.

One possible explanation is that responses in the HAT may be more influenced by short-term emotional states or temporary factors, particularly given the direct involvement of human interaction, an aspect shown to vary depending on handling practices and context, such as movement restrictions or exposure to outdoor environments (Aigueperse & Vasseur, 2021). Previous handling experiences (Waiblinger et al., 2006) could evoke either comfort or fear, while transient factors such as stress, hunger, or fatigue may further impact behaviour during testing. These context-dependent emotional states likely contribute to greater variability in the HAT responses, making them less reflective of stable personality traits compared to tests capturing more generalized behavioural tendencies.

These findings suggest that both tests may reflect a shared underlying trait, likely related to fear (HAT in pigs: level of fear response, Scheffler et al., 2014; SUD in striped mice: fear-anxiety axis, Yuen et al., 2017). However, HAT’s variability across periods may indicate sensitivity to external factors. Further research is therefore needed to determine whether this variability represents a limitation in assessing stable personality traits or if the HAT captures additional context-dependent aspects of reactivity.

#### 5.2.2. The Novel Object Test (NOT), Open Field Test (OFT), and Novel Surface Test (NST)

The inter-test results indicated a lack of consistency across all possible dimension comparisons within the three tests. This aligns with previous research, which has also reported inconsistencies in the personality traits these tests aim to measure. One possible explanation is that behaviours observed in each test may be highly context-dependent. For example, Fox et al. (2009) found no correlation between exploration behaviours in a novel object test and a novel environment test in mountain chickadees, suggesting that exploration in one context does not necessarily translate to another. Similarly, Burns (2008) reported that in guppies, responses in the open field and emergence tests were not correlated with behaviours in the novel object test, calling into question whether the latter is an effective measure of boldness and exploration, traits commonly associated with these tests. In pigs, Haigh et al. (2020) found that the OFT and NOT appeared to measure different aspects of fear or anxiety. While grunting and screaming in the OFT were linked to fear-related behaviours, these vocalizations were associated with longer interaction times in the NOT, suggesting reduced fear in that context.

Conversely, some studies have identified significant relationships between tests. For example, Perals et al. (2017) found that in mynas, the time taken to examine the arena in the OFT was positively linked with the frequency of pecking objects in the NOT, suggesting that exploratory drive was consistently expressed across both contexts. Similarly, Górecka-Bruzda et al. (2011) observed significant positive relationships between behavioural responses (e.g., feeding after exposure to novelty) in the Novel Surface Test (NST) and the static NOT in Polish Cold-blood horses, suggesting that these behaviours may reflect a shared underlying construct.

Given these inconsistencies in the literature, analyzing shared traits across tests remains a challenge. These discrepancies highlight the importance of validating personality assessments across different contexts, as there is still no clear consensus on whether behavioural responses remain stable across the tests used for personality evaluation.

### 5.3. Methodology for Assessing Intra- and Inter-test Consistency

In this study, we addressed the challenge of analysing shared traits across tests by evaluating consistency based on relative rankings rather than absolute behavioural scores. Spearman’s rank correlation was selected as the statistical method for all tests because it accommodates the non-parametric nature of our dataset. This method assigns ranks to individuals and assesses whether these rankings remain relatively stable over time. In other words, while an individual’s absolute score may increase or decrease across periods (e.g., due to habituation or contextual factors), what matters for consistency is whether their position relative to other cows stays similar. For example, a cow that is highly responsive in Period 1 is expected to remain among the most responsive individuals in subsequent periods, even if her exact score or rank changes slightly. This method was preferred over repeatability (R), which measures absolute behavioural consistency, because the focus is on relative consistency compared to conspecifics, rather than requiring the cow to exhibit exactly the same behaviour each time. This aligns with the argument proposed by Stamps & Groothuis (2010), who suggest that studies of animal personality should focus on individual differences relative to others, rather than on absolute behavioural scores.

It is anticipated that behaviours may vary due to environmental factors (e.g., cows may be less active in higher temperatures), but as long as all cows show a similar level of reduced activity, their rankings should remain relatively consistent. This approach is consistent with previous studies such as Zambelis & Vasseur (2022), which used a ranked system to assess divider contact frequency in cows with different stall configurations, evaluating differences in ease of movement. Similar to that study, our focus is not on maintaining the exact behavioural values over time, but rather on ensuring that the same behavioural patterns are observed consistently across testing periods.

Beyond the ranking approach, a possible reason for the lack of significant correlation in the inter-test consistencies could be the unbalanced representation of certain dimensions in the current methodology. For example, the second PCA dimension of the NST might have exhibited an asymmetrical distribution of behaviours. During the naming process of this dimension, the “Activeness” side was supported by behaviours such as running, jumping, placing two hooves on the surface, and zone transitions. However, the “Inactiveness” side did not have any behaviours directly representing it. As a result, it was labeled as the opposite of “Activeness,” based on an interpretive strategy. This inference may have led to overgeneralization, as cows that did not show active behaviours may not necessarily be “inactive.” They could have been engaging in behaviours that were not captured by the PCA and were not inherently inactive (e.g., vocalization, grooming, vigilance, etc.).

Additionally, the scaling of some dimensions does not always follow a simple extreme-to-extreme spectrum. For example, in Dimension 1 of “Reactiveness” in the OFT, the scale ranges from high reactivity (extreme) to low reactivity (neutral), which cannot be directly compared to another dimension, such as Dimension 1 in the NST (Engagement Style), which spans from one extreme (cautious) to another (curiosity). This highlights that comparisons between tests are only valid when the scales are similarly structured. While the dimensions of the tests were created by the researcher and based on subjective interpretations, it is important to recognize that variations in dimension naming and scaling can influence how behaviours are represented and compared across tests. For example, the second dimension of the NST (Interaction Approach), ranging from extreme (activeness) to neutral (inactiveness), could potentially be redefined as activeness to avoidance, providing a different framework for understanding the behaviours. Although dimension labels are subject to the researcher’s interpretation, the absence of supporting behaviours for the ‘inactive’ side could have made this dimension harder to align consistently across tests. This highlights the complexity in measuring behaviours and interpreting dimensions, and although challenges remain in ensuring perfect consistency, even widely accepted frameworks can encounter similar issues. These challenges may stem not only from the analytical method applied, as they reflect the inherent structure of the behavioural tests themselves. Many of these tests are designed to capture extreme responses (e.g., fear, curiosity, boldness), which means intermediate or neutral behaviours are often underrepresented, making PCA-based dimensions uneven or harder to interpret.

Moving forward, ensuring greater variation across the full spectrum of behavioural responses or constructing dimensions based on clearly defined opposing traits may enhance the interpretability of such data. Alternatively, using methods that do not rely on bipolar axes (such as clustering (Amirhosseini et al., 2024), which doesn’t assume binary scales and more effectively captures mixed behavioural signals) may better accommodate dimensions that lack true opposites. These considerations should not devalue the methodology but rather highlight the need for careful attention to dimensional consistency in future work.

## 6. CONCLUSION

The present study aimed to validate five selected behavioural tests by assessing their ability to consistently measure personality traits in dairy cows through repeated testing over time. Of all the tests, only the Sudden Test demonstrated significant consistency across all three time periods, indicating a stable pattern of behavioural expression. In contrast, the remaining tests showed two or fewer significant correlations between periods, suggesting limited temporal reliability. When personality traits were evaluated across different contexts, only the Human Approach Test and Sudden Test showed a moderately strong correlation in two instances. This suggests that these tests may be positively related, as cows showing strong reactions to a sudden stimulus were also more likely to respond intensely to an unknown human. Conversely, no significant inter-test correlations were found among the remaining tests, indicating that they may assess distinct facets of personality or context-specific behaviours. As a result, the Sudden Test emerged as the most promising tool for capturing stable personality traits, while the other tests lacked consistency across both time and context. Future research should focus on standardizing stimuli, minimizing the influence of environmental confounds, and considering longer intervals between tests to further evaluate trait stability. In addition, alternative analytical approaches may help uncover meaningful patterns in behavioural expression that are not solely bidirectional. Together, these improvements could enhance the reliability and practicality of personality assessments, ultimately contributing more targeted and effective welfare interventions in dairy cattle management.

## FUNDING

This project was supported by Natural Sciences and Engineering Research Council of Canada (NSERC Alliance grant; ID# ALLRP 570894-2021), Prompt, Novalait, Dairy Farmers of Canada, Dairy Farmers of Ontario, Les Producteurs de lait du Québec, and Lactanet, through the Vasseur and Diallo Research and Innovation Chair in Animal Welfare and Artificial Intelligence (WELL-E). Additional stipend funding was generously provided by Op+Lait, McGill’s J.W. McConnell Memorial Fellowship, and the McGill Graduate Excellence Fellowship.

## CREDIT AUTHORSHIP CONTRIBUTION STATEMENT

**Xu (Aimee) Xu**: Conceptualization, Data Curation, Formal Analysis, Investigation, Methodology, Visualization, Writing – Original Draft, Writing – Review & Editing. **Marjorie Cellier**: Conceptualization, Formal Analysis, Investigation, Methodology, Supervision, Visualization, Writing – Review & Editing. **Nadège Aigueperse**: Conceptualization, Supervision, Writing – Review & Editing. **Abdoulaye B. Diallo**: Funding Acquisition, Project Administration, Supervision, Writing – Review & Editing. **Elsa Vasseur**: Conceptualization, Funding Acquisition, Methodology, Project Administration, Supervision, Writing – Review & Editing.

## Notes

### Competing Interest Statement

The authors have declared no competing interest.

